# Improved *Cladocopium goreaui* genome assembly reveals features of a facultative coral symbiont and the complex evolutionary history of dinoflagellate genes

**DOI:** 10.1101/2022.07.19.500725

**Authors:** Yibi Chen, Sarah Shah, Katherine E. Dougan, Madeleine J. H. van Oppen, Debashish Bhattacharya, Cheong Xin Chan

## Abstract

Dinoflagellates of the family Symbiodiniaceae are crucial photosymbionts in corals and other marine organisms. Of these algae, *Cladocopium goreaui* is one of the most dominant symbiont species in the Indo-Pacific. Here, we present an improved genome assembly of *C. goreaui* combining new long-read sequence data with earlier generated short-read data. Incorporating new full-length transcripts to guide gene prediction, the *C. goreaui* genome (1.2 Gb) exhibits a high extent of completeness (82.4% based on BUSCO protein recovery) and better resolution of repetitive sequence regions; 45,322 gene models were predicted, and 327 putative, topologically associated domains of the chromosomes were identified. Comparison with other Symbiodiniaceae genomes revealed a prevalence of repeats and duplicated genes in *C. goreaui*, and lineage-specific genes indicating functional innovation. Incorporating 2,841,408 protein sequences from 96 broadly sampled eukaryotes and representative prokaryotes in a phylogenomic approach, we assessed the evolutionary history of *C. goreaui* genes. Of the 5,246 phylogenetic trees inferred from homologous protein sets containing two or more phyla, 35-36% have putatively originated *via* horizontal gene transfer (HGT), predominantly (19-23%) via an ancestral Archaeplastida lineage implicated in the endosymbiotic origin of plastids: 10-11% are of green algal origin, including genes encoding photosynthetic functions. Our results demonstrate the utility of long-read sequence data in resolving structural features of a dinoflagellate genome and highlight how genetic transfer has shaped genome evolution of a facultative symbiont, and more broadly of dinoflagellates.

## Introduction

Dinoflagellates of the family Symbiodiniaceae (LaJeunesse et al., 2018) are diverse microalgae, which are mostly symbionts critical to corals and other coral reef organisms. Symbiodiniaceae provide carbon fixed *via* photosynthesis and other essential nutrients to the coral hosts (Kopp et al., 2013; Muscatine et al., 1984). Environmental stress leads to breakdown of this partnership and loss of the algae, i.e. coral bleaching, putting the corals at risk of starvation, disease, and potential death (Hoegh-Guldberg, 1999). Recent studies of Symbiodiniaceae genomes have revealed extensive sequence and structural divergence (Aranda et al., 2016; Lin et al., 2015; Liu et al., 2018; Shoguchi et al., 2018), and potentially a greater, yet-to-be recognised phylogenetic diversity among these taxa (Dougan et al., 2022; González-Pech et al., 2021). A recent comparative analysis of genomes from 18 dinoflagellate taxa (of which 16 are Symbiodiniaceae) revealed distinct phylogenetic signals between genic and non-genic regions (Lo et al., 2022), indicating differential evolutionary pressures acting on these genomes. This body of research illustrates how the evolutionary complexity of Symbiodiniaceae genomes may explain their diverse symbioses and ecological niches (González-Pech et al., 2019).

*Cladocopium*, the most taxonomically diverse genus in family Symbiodiniaceae, is found predominantly in the Indo-Pacific (Bongaerts et al., 2015; LaJeunesse, 2005), in particular, the species *Cladocopium goreaui* (formally type C1) (Bongaerts et al., 2015; LaJeunesse, 2005). The earlier genome analysis of *C. goreaui* SCF055 (Liu et al., 2018) revealed the genetic capacity of the species to establish and maintain symbiosis with coral hosts, respond to stress, and to undergo meiosis: i.e. many of the implicated genes show evidence of positive selection. Although these results provide insights into the adaptive evolution of genes, the assembled genome, generated using only Illumina short read data, remains fragmented with 41,289 scaffolds (Liu et al., 2018). Additional analysis of the draft genome also indicated that some scaffolds may be of bacterial origin due to their anomalous G+C content (Chen et al., 2020). As such, these data limit our capacity to reliably assess repetitive genomic elements, and evolutionary origins of the predicted genes.

Here we present an improved, hybrid genome assembly for *C. goreaui*, combining novel PacBio long-read sequence data with the existing short-read sequence data from Liu et al. (2018), and incorporating new full-length transcriptome to guide gene prediction. Incorporating proteins predicted from the genome with those from 96 broadly sampled taxa of eukaryotes and representative prokaryotes in a phylogenomic analysis, we also assessed the evolutionary origins of genes in *C. goreaui* and other dinoflagellates, and the impact of horizontal genetic transfer (HGT) in shaping the evolution of this lineage. The earlier investigation based on transcriptome data from the bloom-forming, toxin-producing dinoflagellate *Alexandrium tamarense* (Chan et al., 2012) revealed evidence of HGT, implicating both prokaryote and eukaryote donors, in 14-17% of investigated protein trees. Few genes and no genome data of other dinoflagellates were available when that study was conducted. However, the genomic and genetic resources of dinoflagellates have grown appreciably in the last decade, enabling a more-balanced representation of taxonomic diversity to support such an analysis. In this study we examine how the nuclear genome of *C. goreaui*, and broadly that of dinoflagellates, has evolved and benefited from the acquisition of genomic (and functional) novelties through HGT.

## Results and Discussion

### Improved *C. goreaui* genome assembly reveals more repeats and more duplicated genes

We generated PacBio long-read genome data (50.2 Gbp; Table S1) for *C. goreaui* SCF055 and combined them with existing Illumina short reads in a hybrid approach to generate a *de novo* genome assembly (see Methods). The first published genome assembly of the SCF055 isolate (Liu et al., 2018) was previously refined to exclude putative contaminant sequences (Chen et al., 2020). Compared to the draft assembly reported in Chen et al. (2020), our assembly exhibits a five-fold decrease in the number of scaffolds (6,843) and a three-fold increase in scaffold N50 length (354 Kbp; Table 1). Genome size was estimated at 1.3 Gbp based on *k*-mers (Figure S1), and our improved assembly (1.2 Gbp in size) is larger than the earlier draft (1.0 Gbp; Table 1).

**Table 1.**
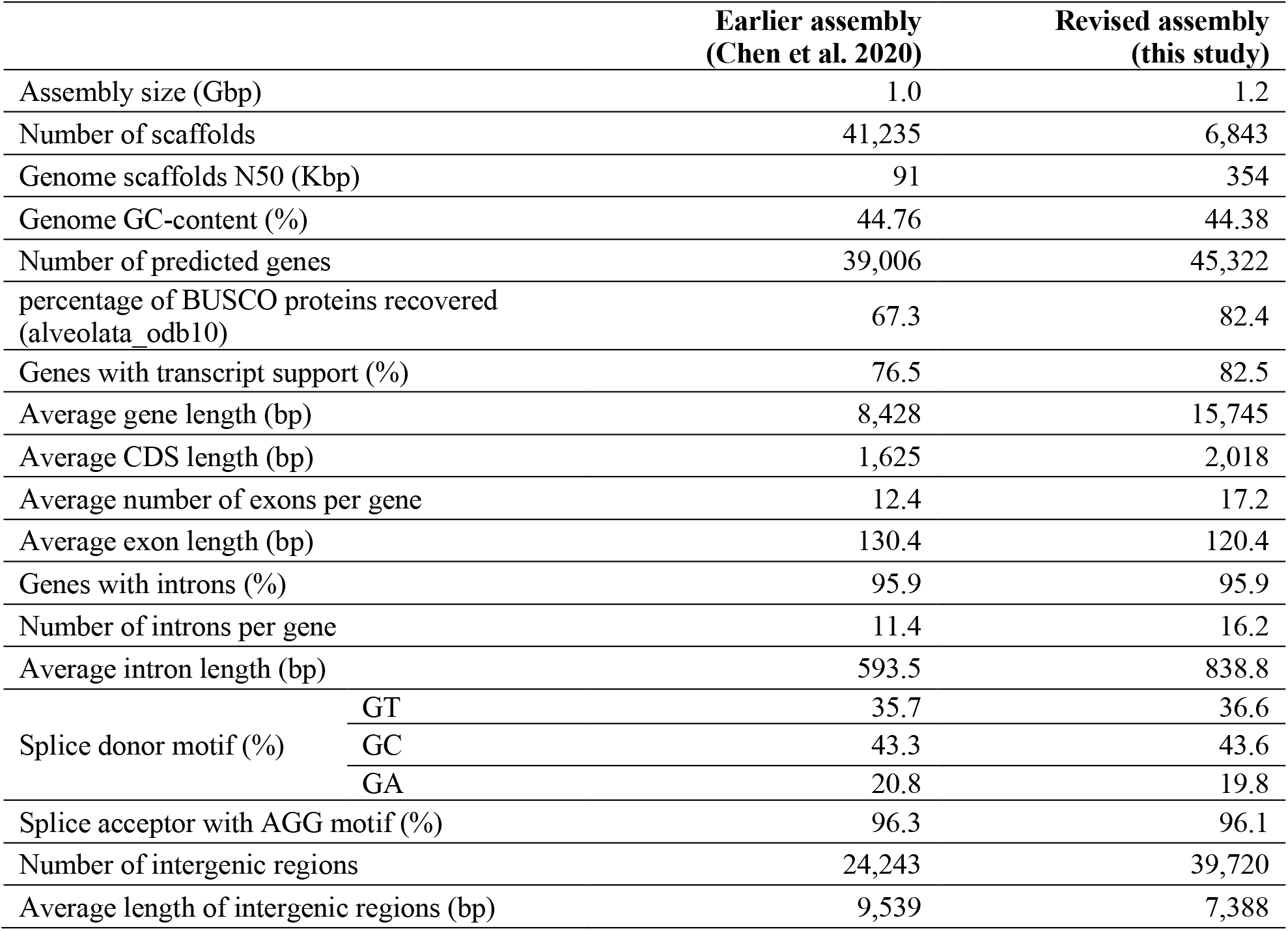
Metrics of the revised genome assembly and predicted genes of *C. goreaui*, compared to the earlier assembled genome.

We also generated 65,432 near full-length transcripts using PacBio Iso-Seq reads to guide prediction of protein-coding genes. Using the same approach tailored for dinoflagellates (Chen et al., 2020), we predicted 45,322 protein-coding genes (mean length 15,745 bp) in the genome, compared to 39,066 (mean length 8,428 bp) reported in Chen et al. (2020). The majority (82.4%) of predicted genes are supported by transcript evidence. These predicted genes are markedly improved, evidenced by the 15.1% greater recovery of core conserved genes (BUSCO alveolate_odb10) (Manni et al., 2021) at 82.4% (Table 1). Most predicted proteins (40,495; 89.3%) have UniProt hits using sequence similarity (BLASTp; *E* ≤ 10^-5^), 19,904 (43.9%) have hits in the curated Swiss-Prot database, 8,836 (19.5%) covering >90% of full-length Swiss-Prot proteins. The remaining 4,827 (10.7%) *C. goreaui* proteins have no significant hits in UniProt, indicating the prevalence of “dark” genes that encode functions yet to be discovered.

We identified and compared the repeat content in the *C. goreaui* genome *versus* the earlier assembly in Chen et al. (2020). Excluding simple repeats, we found a higher repeat content (36.5% of total bases in the assembled genome; Figure 1a) in the current assembly than in the initial data (21.1%), with known repeat types accounting for 17.3% of total bases, compared to 5.6%. This result indicates a better resolution of repetitive regions in the revised genome with the incorporation of long reads. Long terminal repeats (LTR) and DNA transposons are the most prevalent repeat families (constituting 7.3% and 6.2% of total bases, respectively). The two repeat families exhibit distinct levels of sequence divergence: those with Kimura substitution values centred between 0 and 10 are more conserved than those with values centred between 15 and 25. Most LTRs are highly conserved in *C. goreaui*, a trend also observed in the genomes of other dinoflagellates (González-Pech et al., 2021; Stephens et al., 2020).

**Figure 1.**
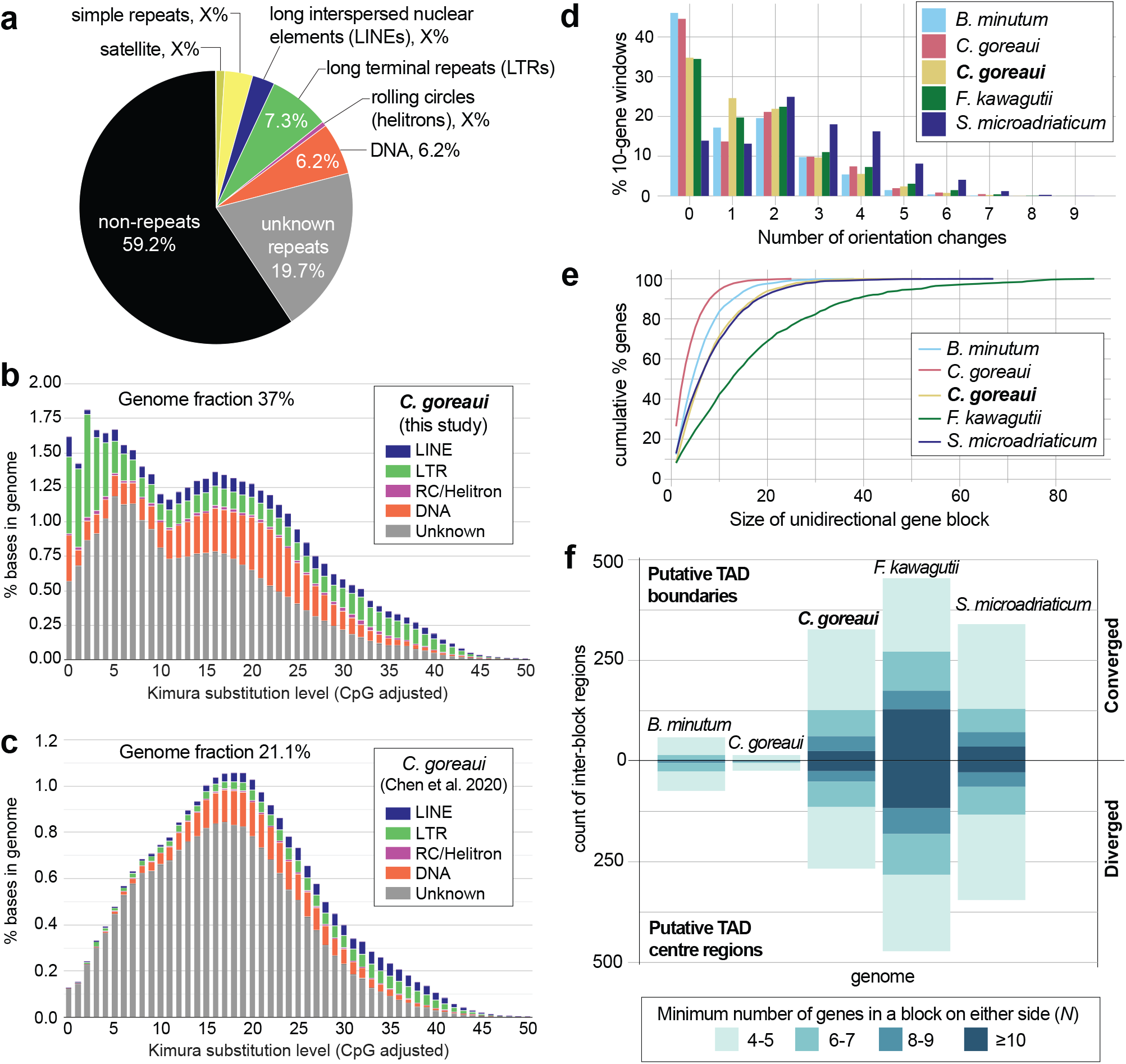
Genome features of *C. goreaui*. (**a**) Repeat families identified in the revised genome assembly; repeat landscape shown for the (**b**) revised genome assembly of *C. goreaui* (highlighted in bold-face) and (**c**) the earlier assembly of from Chen et al. (2020). (**d**) Frequency of strand-orientation change in 10-gene windows and (**e**) cumulative percentage of genes in unidirectionally encoded blocks, shown for five representative Symbiodiniaceae genomes: *Breviolum minutum* (Chen et al. 2020), *C. goreaui* (Chen et al. 2020), *C. goreaui* (boldfaced, this study), *Fugacium kawagutii* (Li et al. 2021), and *Symbiodinium microadriaticum* (Nand et al. 2021); and (**f**) number of inter-block regions in each genome assembly indicating putative TAD central regions and boundaries, shown for representative genomes, based on the minimum number of genes in a unidirectional block. Bars above the *x*-axis represent inter-block regions at which orientations of unidirectional blocks converged, whereas bars below the *x*-axis represent those at which the orientations diverged.

Collinear gene blocks within a genome represent duplicated gene blocks, e.g. *via* segmental or whole-genome duplication. Based on the recovery of these gene blocks, we identified a greater proportion of duplicated genes than Chen et al. (Chen et al., 2020): 35,119 (77.5% of 45,322) genes in duplicates, compared to 25,550 (65.5% of 39,006; Table 2). We found 31,827 (70.2%) genes in dispersed duplicates, suggesting a lack of conserved collinearity of genes in the *C. goreaui* genome. This attribute reflects the extensive structural rearrangements expected in genomes of facultative symbionts (González-Pech et al., 2019). With the improved structural annotation, we recovered a 2.39-fold greater number of tandemly repeated genes, and 387 genes (in 34 collinear blocks) implicated in segmental duplications (Table 2). Tandemly repeated genes in dinoflagellates are thought to be a mechanism for improving transcriptional efficiency, particularly for genes encoding critical functions (Stephens et al., 2020). Comparing Gene Ontology (GO) terms annotated in the 1,998 tandemly repeated genes against those in all *C. goreaui* genes, the top 3 enriched terms for Cellular Component (Table S2) are “chloroplast thylakoid membrane” (GO:0009535; *p* = 1.5 × 10^-7^), “photosystem I reaction center” (GO:0009538; *p* = 5.4 × 10^-7^) and “photosystem II” (GO:0009523; *p* = 0.00042). This result indicates that genes encoding photosynthetic functions tend to appear in tandem repeats in *C. goreaui*, likely to facilitate their transcription. We applied the same approach to genes in segmental duplications and found the most significant enriched terms for this set is Biological Process, “recombinational repair” (GO:0000725, *p* = 7.2 × 10^-5^, Table S3). The repair of errors during genetic recombination is essential for maintaining genome integrity; conservation of collinear organization of genes implicated in this function likely reflects a stronger selective pressure acting on these genes than on the others.

**Table 2.** Types of gene duplication identified in *C. goreaui*.

|  | Earlier assembly<br>(Chen et al. 2020) | Revised assembly<br>(this study) |
| --- | --- | --- |
| Singleton | 13,456 (34.5%) | 10,203 (22.5%) |
| Dispersed | 24,441 (62.6%) | 31,827 (70.2%) |
| Proximal | 273 (0.7%) | 907 (2.0%) |
| Tandem | 836 (2.1%) | 1,998 (4.4%) |
| Segmental | 0 (0.0%) | 387 (0.9%) |

### Topologically associated domains (TADs) and unidirectional gene blocks

Recent studies have clarified interacting genomic regions *via* topologically associated domains (TADs) of dinoflagellate chromosomes (Marinov et al., 2021; Nand et al., 2021). Orientations of unidirectionally encoded gene blocks diverge from a TAD central region, converging at TAD boundaries (Nand et al., 2021). Regulatory elements such as promoters and enhancers for gene expression are hypothesized to be concentrated in these regions to regulate transcription of upstream or downstream unidirectional gene blocks (Lin et al., 2021). To assess putative TAD regions in the revised *C. goreaui* genome, we first identified the unidirectionally encoded gene blocks. We followed Stephens et al. (2020) to enumerate gene-orientation change(s) in a ten-gene window, sliding across entire genome sequences; the tendency for no change in gene orientation is an indication for the prevalence of unidirectional encoding. We performed this analysis in a set of representative Symbiodiniaceae genomes (Figure 1d). Interestingly, we observed a lower percentage (34.7%) of ten-gene windows with conserved orientation in the revised *C. goreaui* genome, when compared to 44.6% in the assembly of Chen et al. (2020). The equivalent percentages in the more-contiguous, near-chromosomal level genome assemblies of *S. microadriaticum* (Nand et al., 2021) (13.9%) and *F. kawagutii* (Li et al., 2020) (34.5%) are also lower, compared to 46.1% in the more-fragmented assembly of *B. minutum* (Chen et al., 2020) (Figure 1d). However, when we assessed the sizes of unidirectional gene blocks in these genomes, they are clearly larger in the more-contiguous assemblies (Figure 1e). For instance, 32.6% of genes in *C. goreaui* are found in block sizes of 10 or more genes, compared to only 7.6% in the earlier assembly (Figure 1e). These observations indicate that the lower recovery of ten-gene windows with conserved orientation is caused by the increased recovery of windows spanning two gene blocks with opposing orientations, as expected in TAD central or boundary regions, in the more-contiguous assemblies. In this way, the more-contiguous assembly enables better recovery of putative TAD regions.

To assess putative TAD regions, we examined genomic regions between any two unidirectional gene blocks. We identified these regions requiring the gene blocks on either side to contain at least *N* number of genes, where *N* is 4, 6, 8, or 10. Figure 1f shows the recovery of these regions across threshold *N* in the representative genomes, with those with converging gene-block orientations (i.e. putative TAD boundaries) above the *x*-axis, and those with diverging orientations (i.e. putative TAD central regions) below the *x*-axis. We recovered orders of magnitude greater numbers of these regions in the more-contiguous *C. goreaui* assembly (e.g. 327 putative TAD boundaries) and in near-chromosomal level assemblies of *S. microadriaticum* (340) and *F. kawagutii* (454), than in the more-fragmented assemblies of *B. minutum* (59) and *C. goreaui* (15). The implicated unidirectional gene blocks on either side of these regions are also larger, e.g. at *N* = 10, we identified 25 putative TAD boundaries in *C. goreaui*, compared to only two in the earlier assembly; the assembly of *F. kawagutii* shows the greatest recovery of TAD-associated regions, with 129 putative TAD boundaries implicating blocks of 10 or more genes on either side. TAD boundaries have been reported to exhibit a dip in GC content in the middle of the sequence (Nand et al., 2021). We observed such a dip in GC content in 17/25 putative TAD boundary regions (at *N* = 10) in *C. goreaui*; an example is shown in Figure S2. Interestingly, our recovery of TAD-associated regions in *C. goreaui* is very similar to that in the chromosome-level assembly of *S. microadriaticum* (Figure 1f), suggesting that our revised assembly, although not derived specifically using chromosome configuration captures, e.g. in Nand et al. (2021), resolves a comparable number of TAD regions.

We also identified genes that tend to disrupt the unidirectional coding of gene blocks, based on their distinct orientation from upstream and downstream genes; such disruptions have been observed in the chromosome-level genome assembly of *S. microadriaticum* (Nand et al., 2021). We identified 3799 (8.4%) of such genes in *C. goreaui*. Interestingly, these genes largely encode functions related to transposon elements. Comparing the annotated GO terms in these genes *versus* those in all *C. goreaui* genes, two most significantly enriched terms for Molecular Function are “nucleic acid binding” (GO:0003676; *p* < 1.0 × 10^-30^) and “RNA-DNA hybrid ribonuclease activity” (GO:0004523; *p* = 1.8 × 10^-20^; Table S4), and the most significantly enriched term for Biological Process is “DNA integration” (GO:0015074; *p* =7.6 × 10^-7^; see Table S4). This result indicates the tendency for mobile genes to disrupt unidirectional coding, and to potentially exploit part of the unidirectional gene block to facilitate transcription.

### Evolutionary origin of *C. goreaui* genes

We inferred a dinoflagellate phylogeny (Supplementary Figure S3) using 3,266 single-copy, strictly orthologous protein sets identified from 1,446,816 sequences of 30 dinoflagellate taxa including *C. goreaui* (Table S5; see Methods). This phylogeny is congruent with the established phylogeny of dinoflagellates (Price & Bhattacharya, 2017; Stephens et al., 2018) with distinct orders occurring in strongly supported clades (bootstrap support [BS] ≥ 90%). *C. goreaui* is placed in a well-supported (BS 100%) clade of family Symbiodiniaceae, and within the order Suessiales (BS 100%) to which the family belongs. This result confirms the phylogenetic position of *C. goreaui* in family Symbiodiniaceae based on putative orthologous proteins, a result that has been demonstrated recently based on whole-genome sequence data using an alignment-free phylogenetic approach (Lo et al., 2022).

We then assessed the evolutionary origin of individual *C. goreaui* genes using protein data. We used 2,841,408 predicted protein sequences from 96 broadly sampled taxa of eukaryotes and prokaryotes (Table S5) to identify 177,346 putative homologous proteins sets based on sequence similarity (see Methods). Of the 45,322 *C. goreaui* proteins, 22,026 (48.6%) are specific to dinoflagellates (i.e. 3,021, 8,748 and 10,257 are specific to *C. goreaui*, order Suessiales, and Dinophyceae; Figure 2a). These results indicate extensive lineage-specific innovation of gene functions following speciation or divergence of dinoflagellates, supporting the notion of extreme divergence of dinoflagellate genes (Dougan et al., 2022; González-Pech et al., 2021; Lin et al., 2015; Stephens et al., 2018).

**Figure 2.**
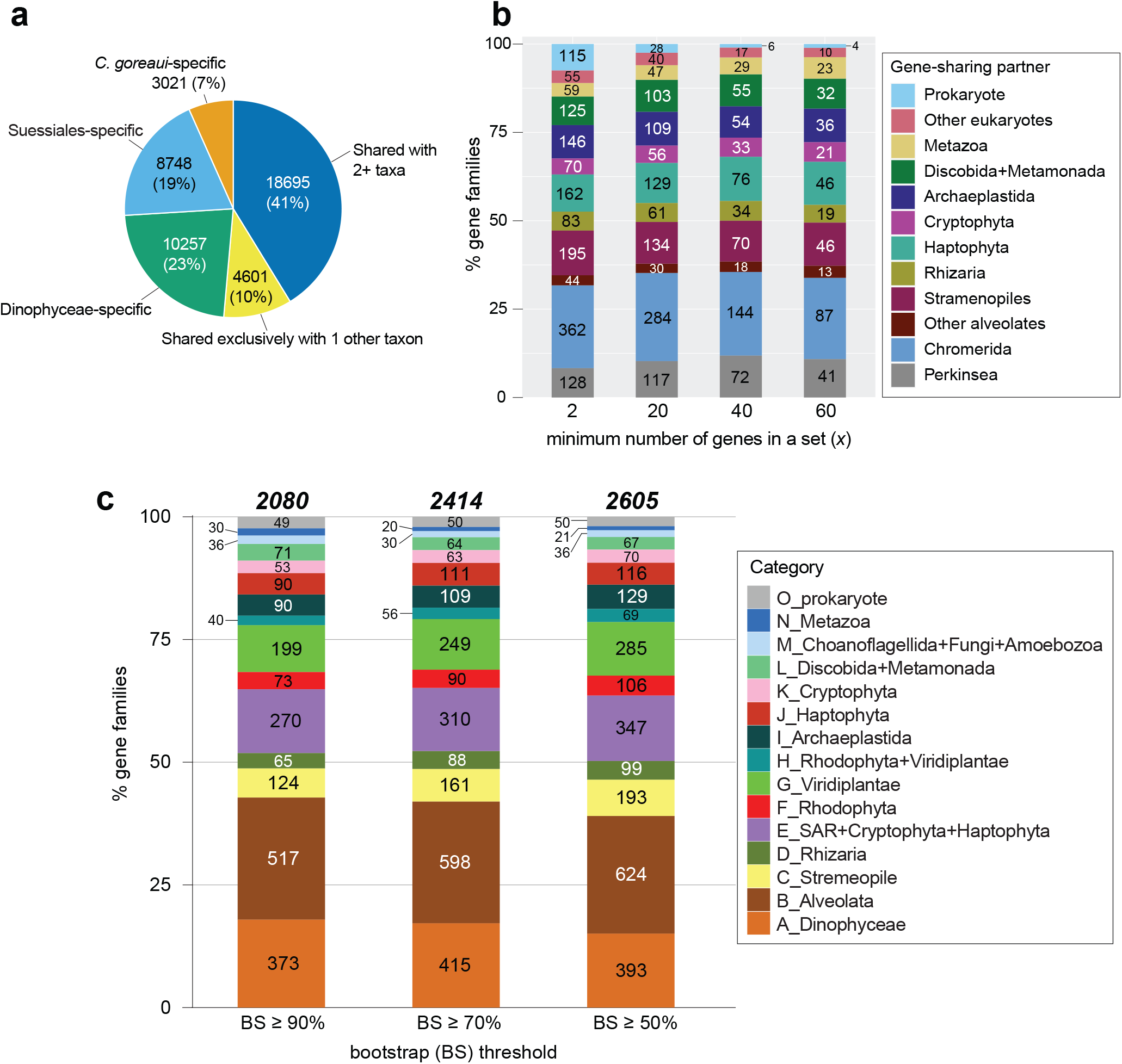
Evolutionary origins of *C. goreaui* genes. (**a**) *C. goreaui* genes classified based on the number of recovered protein homologs in other taxa. (**b**) Distribution of phyla with respect to exclusive gene-sharing partners for *C. goreaui*, based on the number of homologous sets that contain only *C. goreaui* and the other phylum, across the minimum number of genes in each set (*x*) at *x* ≥ 2, ≥ 20, ≥ 40 and ≥ 60. (**c**) Distribution of phyla that are found to share genes with dinoflagellates, based on the number of inferred protein trees in which dinoflagellates and one other phylum were recovered in a monophyletic clade, assessed at bootstrap support (BS) ≥ 90%, ≥ 70% and ≥ 50%.

We found 4,601 (10.2% of 45,322) proteins to be shared exclusively with another phylum, in 1544 homologous sets (Figure 2b). Assuming that inadequate sampling is less of a concern in sets that contain a larger number of genes, we adapted the approach by Chan et al. (2012) to assess these putative gene-sharing partners (phylum) with dinoflagellates, based on the minimum number of genes (*x*) in each set, at *x* ≥ 2, ≥ 20, ≥ 40 and ≥ 60. At *x* ≥ 2, the most frequent gene-sharing partners for dinoflagellates are Chromerida, Perkinsea and other alveolates (534), followed by Stramenopiles (195), Haptophyta (162), and Archaeplastida (146). This result supports the current systematic classification of dinoflagellates in the supergroup Alveolata, the Stramenopiles+Alveolata+Rhizara (SAR) clade (Burki et al., 2020; Hackett et al., 2007), and their close association with haptophytes (Ishida & Green, 2002) and Archaeplastida *via* endosymbiosis implicated by their plastid origin (Chan, Gross, et al., 2011; Yoon et al., 2005). The earlier study based on transcriptome data from the bloom-forming dinoflagellate *Alexandrium tamarense* (Chan et al., 2012) revealed a decrease in proportion of dinoflagellate genes shared with alveolates as *x* increased. This trend is not observed here, e.g. the percentage of genes showing exclusive sharing with alveolates is 34.6%, 37.9%, 38.5% and 37.3% at *x* ≥ 2, ≥ 20, ≥ 40 and ≥ 60. This result suggests that the phylogenetic signal observed here is more robust than in the earlier study, boosted by the greater representation of dinoflagellate taxonomic diversity (30 taxa in Table S5).

The remaining 18,695 (41.2%) *C. goreaui* proteins were recovered in 5,795 homologous sets containing two or more phyla. To assess the evolutionary origins of these genes, we inferred a phylogenetic tree for each of these homologous protein sets; 5,246 remained after passing the initial composition chi-squared test in IQ-TREE to exclude sequences for which the character composition significantly deviates from the average composition in an alignment (see Methods). Among the 5,246 trees, we adopted a computational approach (Stephens et al., 2016) to rapidly identify those in which Dinophyceae taxa form a strongly supported clade with one other phylum, based on observed BS at ≥ 90%, ≥ 70%, and ≥ 50% (Figure 2c; see Methods and Table S6 for detail of our tree-sorting strategy); clades observed at higher BS threshold represent higher confidence results. All sorted trees using the three thresholds are available as Supplementary Data 1. We identified 2,080, 2,414 and 2,605 trees that fit these criteria at BS ≥ 90%, ≥ 70%, and ≥ 50% (Figure 2c); the classification of evolutionary origin for each sorting process is shown in Table S6. The proportions of distinct putative origins for the protein sets are consistent, e.g. those with putative alveolate origin are 24.9%, 24.8% and 24.0% at BS ≥ 90%, ≥ 70%, and ≥ 50% (Figure 2c), reflecting the robust phylogenetic signal we recovered from these data. Remarkably, the evolutionary history of dinoflagellate proteins in more than one-half of the analysed 5,246 trees is too complicated to be classified using our approach, e.g. 2,641 (50.3%) even at our least-stringent threshold of BS ≥ 50%. Some of these proteins (e.g. acyl-CoA dehydrogenase and GTP-binding protein of YchF family) are thought to have a shared origin with fungi or picoprasinophytes (Chan et al., 2012), which are likely to be artefacts due to limited dinoflagellate genome data and sampling bias.

### Genes implicating a history of horizontal transfer

Trees containing a strongly supported monophyletic clade of dinoflagellates and taxonomically remote phyla (Categories F through O in Figure 2c) suggests a history of horizontal genetic transfer; they account for 35.1%, 34.9% and 36.4% of sorted trees at BS ≥ 90%, ≥ 70%, and ≥ 50%. The proportion of HGT-implicated trees is greater than that (14– 17%) in the earlier study based on transcriptome of *Alexandrium tamarense* (Chan et al., 2012). In this study, a more taxonomically balanced set of eukaryote taxa was used, including a larger representation of dinoflagellates and red algae. Therefore, the biases introduced by poor taxon-sampling are diminished, as demonstrated by the robustness of phylogenetic signal that we captured at different stringency levels.

Dinoflagellates possess secondary (and some tertiary) plastids independently acquired from several algal lineages through endosymbioses (Gabrielsen et al., 2011; Yoon et al., 2005). Genes from the endosymbionts were postulated to have been transferred to the host nuclear genome during this process. The implicated endosymbionts include the ancestral Archaeplastida lineages of red and/or green algae, and potentially other eukaryotic microbes e.g. haptophytes, allowing genetic transfer between dinoflagellates and these algal lineages during these events (Ishida & Green, 2002). Secondary plastids found in dinoflagellates and diatoms (stramenopiles) are thought to have arisen from an ancestral red alga (Janouškovec et al., 2010); both red and green algal derived genes have been described in these taxa (Chan, Reyes-Prieto, et al., 2011; Chan et al., 2012; Moustafa et al., 2009).

Here, we focus on the high-confidence trees (clade recovery at BS ≥ 90%) as strong evidence for HGT. Of the 2,080 trees, 402 (19.3%) putatively derived from Archaeplastida (groups F through I in Figure 2c): 73 from Rhodophyta (F), 199 from Viridiplantae (G), and 130 from any combination of Archaeplastida taxa (H and I). At the less-stringent BS threshold, this number is 589 (22.6% of 2,605). *C. goreaui* (and dinoflagellate) genes in these trees likely arose via endosymbiotic genetic transfer due to one or more plastid endosymbioses implicating ancestral Archaeplastida phyla, more evidently with green algae (in Viridiplantae) than with the red algae (Rhodophyta).

Figure 3a shows the phylogenetic tree of beta-glucan synthesis-associated protein homologs, with a strongly supported (BS 97%) monophyletic clade containing Viridiplantae (of green algae *Chlamydomonas, Volvox* and *Chlorella*), haptophytes, Stramenopiles (including diatoms), and dinoflagellates. The beta-glucans are key components of cellulose that form the thecate armour of the cell wall of dinoflagellates (Nevo & Sharon, 1969), and key carbohydrate storage (Salmeán et al., 2017). This tree suggests a putative green algal origin of the gene associated with beta-glucan synthesis in dinoflagellates and other closely related taxa, implicating an ancient HGT among these lineages This is a more parsimonious explanation than to invoke massive gene losses in other alveolates and microbial eukaryotes.

**Figure 3.**
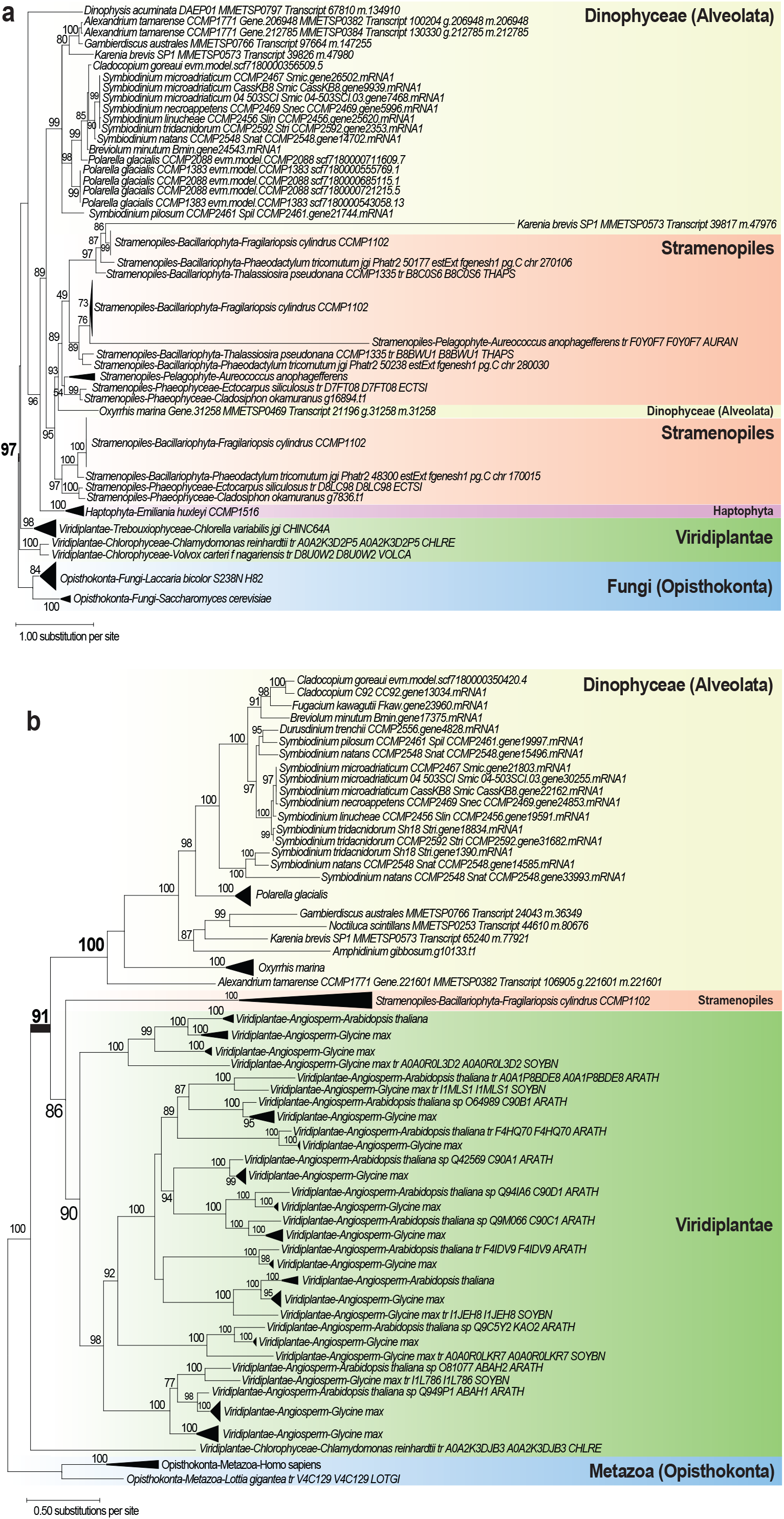
Maximum likelihood trees of (**a**) beta-glucan synthesis-associated protein and (**b**) abscisic acid 8’-hydroxylase, suggesting ancient gene origins from Viridiplantae. The ultrafast bootstrap support of IQ-TREE2 is shown at each internal node; only values ≥70% are shown. Unit of branch length is the number of substitutions per site.

Figure 3b shows the tree for putative abscisic acid 8′-hydroxylase, in which Viridiplantae taxa (largely plants) form a strongly supported (BS 100%) monophyletic clade with the diatom *Fragilariopsis cylindrus* and dinoflagellates. This tree supports a Viridiplantae origin of dinoflagellate genes, and subsequent divergence among the Suessiales (BS 98%) including *C. goreaui*. This enzyme is involved in regulating germination and dormancy of plant seeds *via* oxidation of the hormone abscisic acid (Footitt et al., 2011). It is also known as cytochrome P450 monooxygenase (Krochko et al., 1998). In dinoflagellates, this enzyme is known to regulate encystment and maintenance of dormancy (Deng et al., 2017), and the expression of this gene was found to be upregulated as an initial response to heat stress in a *Cladocopium* species (Rosic et al., 2010). The tree in Figure 3b shows that the protein homologs in dinoflagellates share sequence similarity to those in plants, whereas homologs from other green algae (the more-likely sources of HGT) are absent; given that plants diverged from the green algal lineages, this result suggests that the green algal derived gene in plants, dinoflagellates and the diatom *F. cylindrus* likely subjected to differential functional divergence or gene loss among the green algae.

We also found evidence for more-recent genetic exchanges. Figure 4 shows the tree for a putative sulphate transporter, which has a strongly supported (BS 93%) monophyletic clade containing dinoflagellates and Viridiplantae (mostly green algae), separate from the usual sister lineage of Stremenopiles expected in the SAR grouping in eukaryote tree of life (Burki et al., 2020). This protein is involved in sulphate uptake, which in green algae has direct impact on protein biosynthesis in the plastid (Melis & Chen, 2005). This tree suggests a putative green algal origin of the genes in dinoflagellates that implicates more-recent HGT than those observed in Figure 3. On the other hand, some green algal derived genes appear to have undergone duplication upon diversification of dinoflagellates, e.g. the tree of a hypothetical protein shown in Figure S4. Although the function of this protein in dinoflagellates remains unclear, the homolog in *Arabidopsis thaliana* (UniProt Q94A98; At1g65900) is localised in the chloroplast and implicated in cytokinesis and meiosis (Depuydt & Vandepoele, 2021; Kaundal et al., 2010), lending support to endosymbiotic genetic transfer associated with plastid evolution (Sarai et al., 2020). The fact we found ancient and recent genetic exchanges between green algae and dinoflagellates suggests HGT is a dynamic and ongoing process during genome evolution of dinoflagellates.

**Figure 4.**
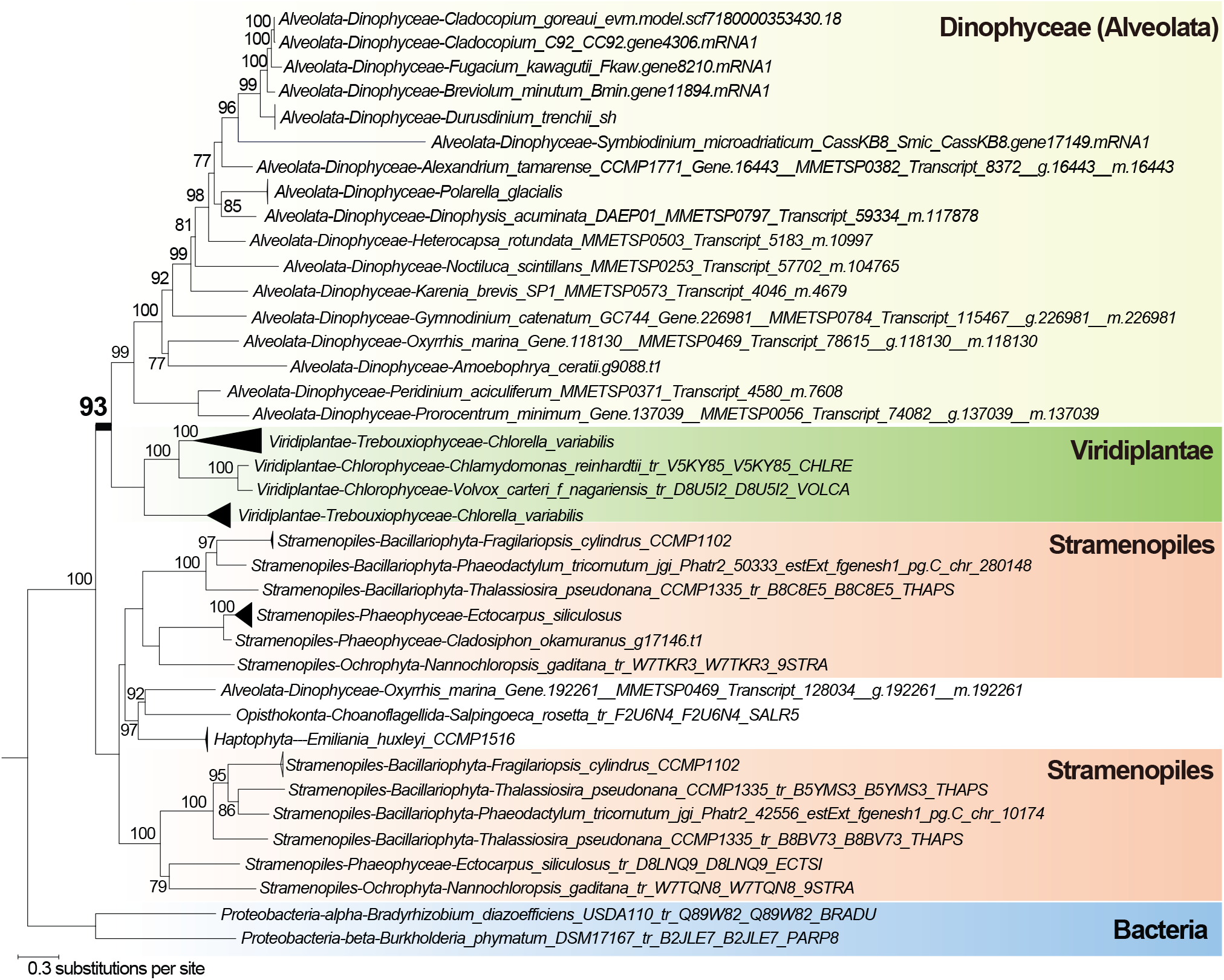
Maximum likelihood trees of putative sulfate transporter, suggesting recent gene origins from Viridiplantae. The ultrafast bootstrap support of IQ-TREE2 is shown at each internal node; only values ≥70% are shown. Unit of branch length is the number of substitutions per site.

Red algal origin of secondary plastids is well established (Janouškovec et al., 2010; Yoon et al., 2002). The stronger signal of green algal than red algal origin we observed here based on a taxonomically broad and dinoflagellate-rich dataset (Table 1) lends support to the notion of an additional cryptic green algal endosymbiosis in the evolution of secondary plastids instead of the “shopping bag” hypothesis that postulates equal proportions of acquired red or green algal genes (Morozov & Galachyants, 2019). Although green algal derived plastids in some dinoflagellates are also known (Archibald & Keeling, 2002; Kamikawa et al., 2015), these taxa are not included in our analysis here due to lack of genome data.

We observed a small proportion (7.1%) of trees that suggest putative genetic exchange between dinoflagellates with distantly related eukaryotes and prokaryotes, e.g. groups L through O in Figure 2c. We cannot dismiss that some of these may be artefacts due to sampling bias or even misidentified sequences in the database. For instance, the phylogeny of phosphatidylinositol 4-phosphate 5-kinase (Figure S5) shows a strongly supported (BS 100%) clade containing 55 dinoflagellate sequences and one sequence from coral *Porites lutea* representing Metazoa; this may be a case of misidentification of the sequence from the dinoflagellate symbiont associated with the coral.

### Genes implicating vertical inheritance

Among the high-confidence trees (recovery at BS ≥ 90%), 64.9% (groups A through E in Figure 2c) provide strong evidence of vertical inheritance; these trees contain a strongly supported (BS ≥ 90%) monophyletic clade containing dinoflagellates only (373), and dinoflagellates plus another closely related taxa of Alveolata (517), Stramenopiles (124), Rhizaria (65), and with SAR group in the presence of Haptophyta and Cryptophyta (270), as expected based on our current understanding of eukaryote tree of life (Burki et al., 2020) Figure S6 shows an example of these trees, specifically, ubiquitin carboxyl-terminal hydrolase. In this tree, all the major phyla are mostly well-resolved in strongly supported clades, e.g. Dinophyceae, Rhizaria, Stramenopiles, and Rhodophyta, each at BS 100%, and the clade of Alveolata+Rhizaria (BS 90%). Figure 5 shows another tree that contains a strongly supported (BS 100%) monophyletic clade of the SAR group, within which three clades (two supported at BS 99%, one at BS 100%), each containing similar dinoflagellate taxa, are recovered, highlighting gene-family expansion. Proteins within the three subclades code for distinct functions: (a) an autophagy-related protein 18a (as with other non-dinoflagellate proteins positioned elsewhere in the tree), (b) a transmembrane protein, and (c) the pentatricopeptide repeat-containing protein GUN1. This observation indicates vertical inheritance of the gene encoding transmembrane protein in dinoflagellates, which was then duplicated and underwent neofunctionalization to generate functional diversity.

**Figure 5.**
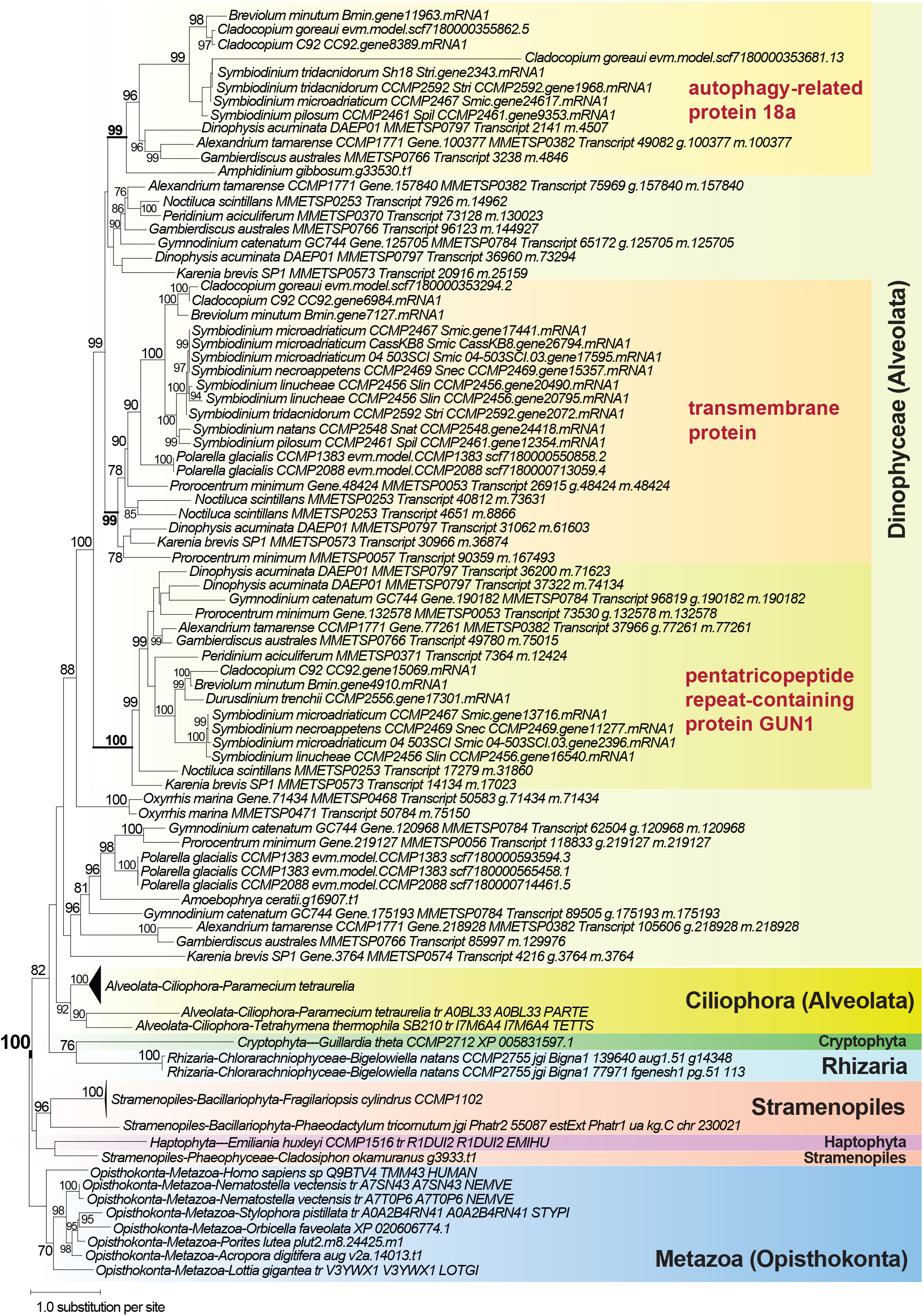
Maximum likelihood protein tree showing vertical inheritance and gene expansion among dinoflagellates, with distinct clades containing a transmembrane protein, the autophagy-related protein 18a, and the pentatricopeptide repeat-containing protein, GUN1. The ultrafast bootstrap support of IQ-TREE2 is shown at each internal node; only values ≥70% are shown. Unit of branch length is the number of substitutions per site.

### Concluding remarks

Our results demonstrate the power of long-read sequence data in elucidating key genome features in *C. goreaui*, at a comparable capacity to chromosome-level genome assemblies of other Symbiodiniaceae, including the resolution of duplicate genes, repetitive genomic elements and TADs. These results support the expected high sequence and structural divergence of dinoflagellate genomes (Dougan et al., 2022; González-Pech et al., 2021). Comparative analysis of the genes revealed clear evidence of lineage-specific innovation in *C. goreaui* and in dinoflagellates generally, implicating about one-half of *C. goreaui* genes; many (52.9%) *C. goreaui* genes remain dark, for which the encoded functions are unknown (Stephens et al., 2018). Our gene-by-gene phylogenetic analysis revealed the intricate evolutionary histories of genes in *C. goreaui* and dinoflagellates, with many too complex to be unambiguously interpreted.

Our results highlight how genetic transfer and gene duplication generated functional diversity and innovation in *C. goreaui*, and in combination with the conserved LTRs and DNA transposons, shaped the genome of this facultative symbiont (González-Pech et al., 2019). The data generated from this study provide a useful reference for future studies of coral symbionts and more broadly of dinoflagellates and microbial eukaryotes. The identified TAD regions, for instance, provides an excellent analysis platform to assess the presence of conserved gene-regulatory elements, e.g. promoters or enhancers of gene expression, as hypothesized in Lin et al. (2021) to facilitate transcription of the unidirectional gene blocks. Our analytic workflow can be adapted and applied to study TADs in other assembled genomes of dinoflagellates.

## Methods

### Generation of long-read genome and transcriptome data

*Cladocopium goreaui* SCF055-01 is a single-cell monoclonal culture first isolated from the coral *Acropora tenuis* at Magnetic Island, Queensland, Australia (Howells et al., 2012). The cultures were maintained at the Australian Centre of Marine Science (AIMS) in Daigo’s IMK medium at 26°C, 90 uE/cm^2^/s^-1^. High molecular weight genomic DNA was extracted following the SDS method described in Wilson et al. (2002).The sample was sent to Ramaciotti Centre for Genomics (University of New South Wales, Sydney) for sequencing using the PacBio, first using RS II, then the Sequel platform (Table S1). DNA fragments of lengths 10–20 Kb were selected for the preparation of sequencing libraries. In total, 6.2 million subreads were produced (50.2 Gbp).

Total RNA was extracted from cultured SCF055 cells in exponential growth phase (∼10^6^ cells), combining the standard Trizol method with Qiagen RNeasy protocol, following Rosic and Hoegh-Guldberg (Rosic & Hoegh-Guldberg, 2010). Quality and quantity of RNA were assessed using a Bioanalyzer and Qubit. The RNA sample was sent to the sequencing facility at the University of Queensland’s Institute for Molecular Bioscience for generation of Iso-Seq data using the PacBio Sequel II platform. Iso-Seq library was prepared using the NEBNext Single Cell/Low Input cDNA Synthesis and Amplification Module and the SMRTbell Express Template Prep Kit 2.0 following standard protocol. Sequencing was conducted using half of a Sequel II SMRT cell. From these raw data, we generated 3,534,837 circular consensus sequencing (CCS) reads (7.3 Gb; average 36 passes) using CCS v4.2.0. The CCS reads were then fed into the Iso-Seq pipeline v3.3.0 pipeline for standard Iso-Seq processing, which includes key steps of read refinement, isoform clustering, and polishing, resulting in 55,505 high-quality, non-redundant, polished Iso-Seq transcripts (total 79 Mb; N50 length 1,493 bases).

### *De novo* genome assembly combining short- and long-read sequence data

We combined the long-read sequence data with all short-read sequence data from Liu et al. (2018) in a hybrid genome assembly using MaSuRCA v3.4.2 (Zimin et al., 2017). Because the SCF055 culture is xenic, we adapted the approach in Iha et al. (2021) to identify and remove putative contaminant sequences from bacterial, archaeal, and viral sources. Bowtie2 (Langmead & Salzberg, 2012) was first used to map the genome sequencing data (Illumina paired-end reads) using the --very-fast algorithm to the assembled genome scaffolds to obtain information of sequencing depth. BlobTools v1.1 (Laetsch & Blaxter, 2017) was then used to identify anomalies of GC content and sequencing depth among the scaffolds, and to assign a taxon to each scaffold (using the default “bestsum” algorithm) based on hits in a BLASTn (*E* ≤ 10^-20^) search against the NCBI nucleotide (nt) database release 2020-01-08. We also mapped available transcriptome data onto the assembled genome to further assess gene structure to aid identification of intron-containing genes on the scaffolds as indication of eukaryote origin. To do this, we used our Iso-Seq transcripts (above) and the RNA-Seq data from Levin et al. (2016) that we assembled using Trinity v2.9.1 (Grabherr et al., 2011) in *de novo* mode. Mapping was conducted using minimap2 v2.17-r975-dirty (Li, 2018) (--secondary=no -ax splice:hq -uf –splice-flank=no), for which the code has been modified to recognise alternative splice-sites in dinoflagellate genes. Using the taxon assignment, genome covarege, and transcript support information, we identified and removed putative contaminant sequences from the genome assembly following a decision tree based on these results (Figure S7).

### Estimation of genome size based on sequencing data

To estimate the genome size, we adapted the *k*-mer-based approach used by González-Pech et al. (2021). We first enumerated *k*-mers of size *k* = 21 from the sequencing reads using Jellyfish v2.3.0 (Marçais & Kingsford, 2011). The resulting histogram (jellyfish histo -- high=1000000) of k-mer count was used as input for GenomeScope2 (Ranallo-Benavidez et al., 2020) to estimate a haploid genome size. Genome of C. goreaui (and other Symbiodiniaceae) are thought to be haploid (Liu et al., 2018) and we did not observe bimodal distribution of *k*-mer coverage expected in a diploid genome (Figure S1).

### Annotation of repeat content

*De novo* repeat families were predicted from the genome assembly using RepeatModeler v1.0.11 (http://www.repeatmasker.org/RepeatModeler/). All repeats (including known repeats in RepeatMasker database release 20181026) were identified and masked using RepeatMasker v4.0.7 (http://www.repeatmasker.org/); the masked sequences were used for *ab initio* gene prediction (below). The repeat landscape plot was generated with Perl script *createRepeatLandscape*.*pl* available from RepeatMasker.

### *Ab initio* prediction of protein-coding genes

To predict protein-coding genes from the assembled genome sequences, we adopted the approach in Chen et al. (2020), using the workflow tailored for dinoflagellate genomes (https://github.com/TimothyStephens/Dinoflagellate_Annotation_Workflow), which was also applied in earlier studies of Symbiodiniaceae genomes (González-Pech et al., 2021; Liu et al., 2018)

Briefly, the transcriptome data (combining our 55,505 high-quality Iso-Seq transcripts and data from Levin et al. (2016); above) were mapped onto the assembled genome with Minimap2 (Li, 2018). All transcripts were combined into gene assemblies using PASA v2.3.3 (Haas et al., 2003) for which the code was modified to recognise alternative splice sites (available at https://github.com/chancx/dinoflag-alt-splice). TransDecoder v5.2.0 (Haas et al., 2003) was used to predict open reading frames on the PASA-assembled transcripts; these represent the transcript-based predicted genes. The predicted proteins were searched (BLASTP, *E* ≤ 10^-20^, >80% query cover) against a protein database consisting of RefSeq proteins (release 88) and predicted proteins of available Symbiodiniaceae genomes (Table S7). The gene models were checked for transposable elements using HHblits v2.0.16 (Remmert et al., 2012) and TransposonPSI (http://transposonpsi.sourceforge.net/), searching against the JAMg transposon database (https://github.com/genomecuration/JAMg); those containing these elements were removed from subsequent steps. After removal of redundant sequences based on similarity using CD-HIT v4.6.8 (Li & Godzik, 2006) (-c 0.75 -n 5), the final curated gene models were used to identify high-quality “golden genes” using the script *Prepare_golden_genes_for_predictors*.*pl* from the JAMg pipeline (http://jamg.sourceforge.net/), altered to recognise alternative splice sites.

We used four other programs for predicting protein-coding genes. The “golden genes” above were used as a training set for SNAP (Korf, 2004) and AUGUSTUS v3.3.1 (Stanke et al., 2006) to predict genes from the repeat-masked genome; the code for AUGUSTUS was altered to recognise alternative splice sites of dinoflagellates (available at https://github.com/chancx/dinoflag-alt-splice). We also used GeneMark-ES (Lomsadze et al., 2005) and MAKER v2.31.10 (Holt & Yandell, 2011) for which code was modified to recognise alternative splice sites, in *protein2genome* mode guided by SwissProt database (retrieved 27 June 2018) and other predicted proteins from Symbiodiniaceae (Table S7). Finally, gene predictions from all five methods, i.e. the *ab initio* predictions (from GeneMark-ES, AUGUSTUS, and SNAP), MAKER protein-based predictions, and PASA transcript-based predictions, were integrated using EvidenceModeler v1.1.1 (Haas et al., 2008) to yield the gene models; see Chen et al. (2020) for detail. The gene models were further polished with PASA (Haas et al., 2003) for three rounds to incorporate isoforms and UTRs, yielding the final gene models.

### Functional annotation of *C. goreaui* genes

For functional annotation, all predicted proteins were searched against all protein sequences on Uniport (release 2021_03). Only hits with *E* ≤ 10^-5^ were retained. Gene Ontology (GO) terms associated with top hits were first retrieved from Uniprot website using the “Retrieve/ID mapping tool”, then mapped to the corresponding queries.

### Analysis of duplicated genes

To identify and classify duplicated genes, we first performed all-versus-all BLASTp (*E* ≤ 10^-^ 5) on corresponding proteins of all predicted genes. The top five hits (excluding the query itself) were used as input for MCScanX (Wang et al., 2013) in *duplicate_gene_classifier* mode to classify genes into five categories: singleton (single-copy genes), dispersed (paralogs away from each other; i.e. at least 20 genes apart), proximal (paralogs near each other), tandem (paralogs in tandem gene block) and segmental (duplicates of collinear blocks).

### GO enrichment analysis

R package topGO (Alexa & Rahnenführer, 2009) was used for enrichment analysis of GO terms. In total, 21,356 genes were annotated with one or more GO terms; these were used as the background set. Genes in tandem repeats and segmental duplication are used as the test set to search for enriched GO terms, in independent analyses. Fisher’s exact test is applied to assess statistical significance, instances with p ≤ 0.01 is considered significant.

### Analysis of unidirectional gene blocks and TADs

For this part of analysis, we focused on five representative assembled genomes of dinoflagellates: *C. goreaui* from this study, *C. goreaui* from Chen et al. (Chen et al., 2020)*Fugacium Kawagutii* (Li et al., 2020), *Breviolum minutum* (Chen et al., 2020), and *Symbiodinium microadriaticum* (Nand et al., 2021). Unidirectional gene blocks were identified based on a block of genes within which their orientiations are the same. A putative TAD boundary is the region at which the orientations of two blocks converged. A putative TAD central region is the region at which the orientations diverged. We analysed putative TAD regions and unidirectional gene blocks based on the minimum number of genes within a block, *N*, at *N* = 4, 6, 8 and 10.

To validate the putative TADs, we searched for GC dip in the TAD boundaries, following Nand et al. (Nand et al., 2021). On each scaffold, for each sliding 4Kb-window, we calculated the localised G+C content. A putative region of GC dip is identified based on three criteria: (1) the G+C in a 4Kb region is lower than average GC content of the entire scaffold (i.e. the background); (2) the largest difference between the localised G+C and the background G+C is larger than 0.05%; and that (3) the implicated region is longer than 5000 bp.

### Phylogenomic analysis of *C. goreaui* genes

To investigate the putative origins of *C. goreaui* genes, we compiled a comprehensive protein database (2,841,408 sequences) of 96 broadly sampled taxa from diverse lineages, encompassing eukaryotic, bacterial, and archaeal taxa; of which 30 are dinoflagellates (Table S5). For species where there were multiple datasets for the same isolate, the protein sequences were clustered at 90% sequence identity using CD-HIT-v4.8.1(Li & Godzik, 2006) to reduce redundancy. Isoforms are reduced to retain one representative protein (longest) per genes.

Using all 2,841,408 protein sequences from the database (Table S5), homologous protein sets were inferred using OrthoFinder-v2.3.8 (Emms & Kelly, 2015) at default setting. For this analysis, we restricted our analysis to 177,346 putative homologous sets of *C. goreaui* proteins (i.e. sets in which one or more *C. goreaui* sequence is represented). For homologous sets contain only Dinophyceae and one other phylum (an exclusive gene-sharing partner), the putative gene-sharing partner was assessed based on the number of implicated homologous sets, requiring at least *x* number of genes in each set (for *x* = 2, 20, 40, and 60). For the other protein sets, multiple sequence alignment was performed using MAFFT-v7.453 (Katoh & Standley, 2013) with parameters *--maxiterate 1000 --localpair*. Following Stephens et al. 2018 (Stephens et al., 2018), ambiguous and non-phylogenetically informative sites in each alignment were trimmed using trimAl-v1.4.1 (Capella-Gutiérrez et al., 2009) in two steps: trimming directly with *-automated1*, then with *-resoverlap 0*.*5 -seqoverlap 50*. A maximum likelihood tree for each protein set was inferred from these trimmed alignments, using IQ-TREE2 (Minh et al., 2020), with an edge-unlinked partition model and 2000 ultrafast bootstraps. The initial step of IQ-TREE2 by default is to perform a composition chi-squared test for every sequence in an alignment, the sequence for which the character composition significantly deviates from the average composition in the alignment is removed. Alignments filtered this way were further removed if the target *C. goreaui* sequence was removed, and if the alignment contains no more than four sequences. In total, 5,246 trees were used in subsequent analysis.

### Inference of the dinoflagellate species tree

To infer the dinoflagellate species tree, we first inferred homologous protein sets with OrthoFinder-v2.3.8 for the 30 dinoflagellate taxa and *Perkinsus marinus* (outgroup) in Table S5. The 3,266 strictly orthologous, single-copy protein sets (i.e. sets in which each dinoflagellate taxon is represented no more than once) were used for inferring the species tree. For each set, multiple sequence alignment was performed, the alignment was trimmed, per our approach described above. A consensus maximum likelihood reference species tree was then inferred using IQ-TREE2, with an edge-unlinked partition model and 2000 ultrafast bootstraps.

### Inference of *C. goreaui* gene origins

Putative origins of *C. goreaui* genes were identified based on the presence of strongly supported clades (determined at a bootstrap support threshold) that include *C. goreaui* (and/or other dinoflagellates) and one other taxon group (e.g., a phylum). We used PhySortR (Stephens et al., 2016) to quickly sort through thousands of protein trees (i.e. we assume these as gene trees) for the specific target clades, independently at bootstrap thresholds of ≥90% (more stringent, higher confidence), ≥70%, and ≥50% (less stringent, lower confidence); default values were used for other parameters. Our 176-step tree-sorting strategy is detailed in Table S6. Briefly, we sorted the trees based on recovery of a strongly supported monophyletic clade containing both the subject group (dinoflagellates) and target group in three stages: first (a) with target groups implicated in endosymbiosis in the evolution of plastid (e.g. Archaeplastida); then (b) with closely related target group expected under vertical inheritance, and finally (c) with other remotely related eukaryote or prokaryote target group as further indication of horizontal genetic transfer. At each stage, the subject group would proceed from the most inclusive (i.e. dinoflagellates plus other closely related taxa in the SAR supergroup, Cryoptophyta and Haptophyta), and progressively the most distant lineage was removed in each iterative sorting, leaving only dinoflagellates. For each subject group, the target group would proceed similarly from the most inclusive, e.g. in stage one, all three phyla of Archaeplastida, to subsequent individual phylum.

## Supplementary Materials

**Figure S1**. Genome size estimation for *Cladocopium goreaui* using GenomeScope v2.0.

**Figure S2**. An example of G+C dip observed in the *C. goreaui* genome of a putative boundary of topologically associated domain (TAD).

**Figure S3**. Species tree of 29 dinoflagellate taxa and *Perkinsus* as outgroup, inferred from 3,266 strictly orthologous (single-copy) protein sets.

**Figure S4**. Maximum likelihood tree showing gene expansion of a green algal derived gene family.

**Figure S5**. Maximum likelihood tree of phosphatidylinositol 4-phosphate 5-kinase showing possible misidentification of the sequence from the dinoflagellate symbiont associated with the coral.

**Figure S6**. Maximum likelihood tree of ubiquitin carboxyl-terminal hydrolase showing strong evidence of vertical inheritance.

**Figure S7**. Decision tree for identification and removal of putative contaminant sequences.

**Table S1**. PacBio long-read genome sequencing data from *C. goreaui* generated in this study.

**Table S2**. Enriched GO terms for tandemly repeated genes, shown for Biological Process (BP), Cellular Component (CC), and Molecular Function (MF).

**Table S3**. Enriched GO terms for genes in segmental duplicates, shown for Biological Process (BP), Cellular Component (CC), and Molecular Function (MF).

**Table S4**. Enriched GO terms for genes disrupting unidirectional coding of gene blocks, shown for Biological Process (BP), Cellular Component (CC), and Molecular Function (MF).

**Table S5**. Protein database for phylogenomic analysis.

**Table S6**. Sorting order of phylogenetic trees and the associated classifications.

**Table S7**. Protein sequences used to guide *ab initio* prediction of protein-coding genes.

**Data S1**. All sorted trees where Dinophyceae taxa form a strongly supported clade with one other phylum

## Author Contributions

Conceptualization, Y.C., D.B. and C.X.C.; Methodology, Y.C., S.S., K.E.D. and C.X.C.; Formal analysis, Y.C. and S.S.; Writing – Original Draft Preparation, Y.C.; Writing – Review and Editing, all authors; Supervision, C.X.C.; Funding Acquisition, C.X.C. and D.B.

## Funding

This research was funded by Australian Research Council grants DP150101875 and DP190102474 (C.X.C. and D.B.), and the University of Queensland Research Training Program (Y.C. and S.S.). M.J.H.v.O. acknowledges Australian Research Council Laureate Fellowship FL180100036.

## Data Availability Statement

Genome and transcriptome long-read sequence data generated from this study are available at NCBI Sequence Read Archive (BioProject accession XXXXXXXXXX). The assembled genome, predicted gene models and proteins from this study are available at https://cloudstor.aarnet.edu.au/plus/s/ghXS5Qtf3b4eZA4.

## Acknowledgements

This project is supported by high-performance computing facilities the National Computational Infrastructure (NCI) National Facility systems through the NCI Merit Allocation Scheme (Project d85) awarded to CXC, the University of Queensland Research Computing Centre, and HPC from the Australian Centre for Ecogenomics at UQ. We are grateful to technical assistance provided by Carlos Alvarez Roa and Lesa Peplow in extractions of DNA and RNA from *C. goreaui* SCF055 at the Symbiont Culturing Facility at the Australian Institute of Marine Science in Townsville, Queensland.

## Conflicts of Interest

The authors declare no conflict of interest. The funders had no role in the design of the study; in the collection, analyses, or interpretation of data; in the writing of the manuscript; or in the decision to publish the results.

## Notes

https://cloudstor.aarnet.edu.au/plus/s/ghXS5Qtf3b4eZA4

### Competing Interest Statement

The authors have declared no competing interest.

## References

Alexa, A., & Rahnenführer, J. (2009). Gene set enrichment analysis with topGO. Bioconductor Improv, 27, 1–26.

Aranda, M., Li, Y., Liew, Y. J., Baumgarten, S., Simakov, O., Wilson, M. C., Piel, J., Ashoor, H., Bougouffa, S., & Bajic, V. B. (2016). Genomes of coral dinoflagellate symbionts highlight evolutionary adaptations conducive to a symbiotic lifestyle. Scientific Reports, 6, 39734.

Archibald, J. M., & Keeling, P. J. (2002). Recycled plastids: a ‘green movement’ in eukaryotic evolution. Trends in Genetics, 18(11), 577–584. https://doi.org/10.1016/s0168-9525(02)02777-4

Bongaerts, P., Carmichael, M., Hay, K. B., Tonk, L., Frade, P. R., & Hoegh-Guldberg, O. (2015). Prevalent endosymbiont zonation shapes the depth distributions of scleractinian coral species. Royal Society open science, 2(2), 140297.

Burki, F., Roger, A. J., Brown, M. W., & Simpson, A. G. B. (2020). The New Tree of Eukaryotes. Trends in Ecology & Evolution, 35(1), 43–55. https://doi.org/10.1016/j.tree.2019.08.008

Capella-Gutiérrez, S., Silla-Martínez, J. M., & Gabaldón, T. (2009). trimAl: a tool for automated alignment trimming in large-scale phylogenetic analyses. Bioinformatics, 25(15), 1972–1973.

Chan, C. X., Gross, J., Yoon, H. S., & Bhattacharya, D. (2011). Plastid origin and evolution: new models provide insights into old problems. Plant Physiology, 155(4), 1552–1560. https://doi.org/10.1104/pp.111.173500

Chan, C. X., Reyes-Prieto, A., & Bhattacharya, D. (2011). Red and green algal origin of diatom membrane transporters: insights into environmental adaptation and cell evolution. PLoS ONE, 6(12), e29138. https://doi.org/10.1371/journal.pone.0029138

Chan, C. X., Soares, M. B., Bonaldo, M. F., Wisecaver, J. H., Hackett, J. D., Anderson, D. M., Erdner, D. L., & Bhattacharya, D. (2012). Analysis of Alexandrium tamarense (Dinophyceae) genes reveals the complex evolutionary history of a microbial eukaryote Journal of Phycology, 48(5), 1130–1142.

Chen, Y., González-Pech, R. A., Stephens, T. G., Bhattacharya, D., & Chan, C. X. (2020). Evidence that inconsistent gene prediction can mislead analysis of dinoflagellate genomes. Journal of Phycology, 56(1), 6–10.

Deng, Y., Hu, Z., Shang, L., Peng, Q., & Tang, Y. Z. (2017). Transcriptomic Analyses of Scrippsiella trochoidea Reveals Processes Regulating Encystment and Dormancy in the Life Cycle of a Dinoflagellate, with a Particular Attention to the Role of Abscisic Acid. Frontiers in Microbiology, 8. https://doi.org/10.3389/fmicb.2017.02450

Depuydt, T., & Vandepoele, K. (2021). Multi-omics network-based functional annotation of unknown Arabidopsis genes. The Plant Journal, 108(4), 1193–1212. https://doi.org/https://doi.org/10.1111/tpj.15507

Dougan, K. E., González-Pech, R. A., Stephens, T. G., Shah, S., Chen, Y., Ragan, M. A., Bhattacharya, D., & Chan, C. X. (2022). Genome-powered classification of microbial eukaryotes: focus on coral algal symbionts. Trends in Microbiology. https://doi.org/10.1016/j.tim.2022.02.001

Emms, D. M., & Kelly, S. (2015). OrthoFinder: solving fundamental biases in whole genome comparisons dramatically improves orthogroup inference accuracy. Genome Biology, 16(1), 157.

Footitt, S., Douterelo-Soler, I., Clay, H., & Finch-Savage, W. E. (2011). Dormancy cycling in Arabidopsis seeds is controlled by seasonally distinct hormone-signaling pathways. Proceedings of the National Academy of Sciences, 108(50), 20236–20241. https://doi.org/10.1073/pnas.1116325108

Gabrielsen, T. M., Minge, M. A., Espelund, M., Tooming-Klunderud, A., Patil, V., Nederbragt, A. J., Otis, C., Turmel, M., Shalchian-Tabrizi, K., & Lemieux, C. (2011). Genome evolution of a tertiary dinoflagellate plastid. PLoS ONE, 6(4), e19132.

González-Pech, R. A., Bhattacharya, D., Ragan, M. A., & Chan, C. X. (2019). Genome evolution of coral reef symbionts as intracellular residents. Trends in Ecology & Evolution, 34(9), 799–806.

González-Pech, R. A., Stephens, T. G., Chen, Y., Mohamed, A. R., Cheng, Y., Shah, S., Dougan, K. E., Fortuin, M. D. A., Lagorce, R., Burt, D. W., Bhattacharya, D., Ragan, M. A., & Chan, C. X. (2021). Comparison of 15 dinoflagellate genomes reveals extensive sequence and structural divergence in family Symbiodiniaceae and genus Symbiodinium. BMC Biology, 19(1), 73. https://doi.org/10.1186/s12915-021-00994-6

Grabherr, M. G., Haas, B. J., Yassour, M., Levin, J. Z., Thompson, D. A., Amit, I., Adiconis, X., Fan, L., Raychowdhury, R., & Zeng, Q. (2011). Trinity: reconstructing a full-length transcriptome without a genome from RNA-Seq data. Nature Biotechnology, 29(7), 644.

Haas, B. J., Delcher, A. L., Mount, S. M., Wortman, J. R., Smith Jr, R. K., Hannick, L. I., Maiti, R., Ronning, C. M., Rusch, D. B., & Town, C. D. (2003). Improving the Arabidopsis genome annotation using maximal transcript alignment assemblies. Nucleic Acids Research, 31(19), 5654–5666. https://doi.org/10.1093/nar/gkg770

Haas, B. J., Salzberg, S. L., Zhu, W., Pertea, M., Allen, J. E., Orvis, J., White, O., Buell, C. R., & Wortman, J. R. (2008). Automated eukaryotic gene structure annotation using EVidenceModeler and the Program to Assemble Spliced Alignments. Genome Biology, 9(1), R7. https://doi.org/10.1186/gb-2008-9-1-r7

Hackett, J. D., Yoon, H. S., Li, S., Reyes-Prieto, A., Rümmele, S. E., & Bhattacharya, D. (2007). Phylogenomic analysis supports the monophyly of cryptophytes and haptophytes and the association of rhizaria with chromalveolates. Molecular Biology and Evolution, 24(8), 1702–1713. https://doi.org/10.1093/molbev/msm089

Hoegh-Guldberg, O. (1999). Climate change, coral bleaching and the future of the world’s coral reefs. Marine and Freshwater Research, 50(8), 839–866.

Holt, C., & Yandell, M. (2011). MAKER2: an annotation pipeline and genome-database management tool for second-generation genome projects. BMC Bioinformatics, 12(1), 491. https://doi.org/10.1186/1471-2105-12-491

Howells, E., Beltran, V., Larsen, N., Bay, L., Willis, B., & Van Oppen, M. (2012). Coral thermal tolerance shaped by local adaptation of photosymbionts. Nature Climate Change, 2(2), 116–120.

Iha, C., Dougan, K. E., Varela, J. A., Avila, V., Jackson, C. J., Bogaert, K. A., Chen, Y., Judd, L. M., Wick, R., & Holt, K. E. (2021). Genomic adaptations to an endolithic lifestyle in the coral-associated alga Ostreobium. Current Biology, 31(7), 1393–1402. e1395.

Ishida, K.-i., & Green, B. R. (2002). Second-and third-hand chloroplasts in dinoflagellates: Phylogeny of oxygen-evolving enhancer 1 (PsbO) protein reveals replacement of a nuclear-encoded plastid gene by that of a haptophyte tertiary endosymbiont. Proceedings of the National Academy of Sciences, 99(14), 9294–9299. https://doi.org/10.1073/pnas.142091799

Janouškovec, J., Horák, A., Oborník, M., Lukeš, J., & Keeling, P. J. (2010). A common red algal origin of the apicomplexan, dinoflagellate, and heterokont plastids. Proceedings of the National Academy of Sciences, 107(24), 10949–10954. https://doi.org/doi:10.1073/pnas.1003335107

Kamikawa, R., Tanifuji, G., Kawachi, M., Miyashita, H., Hashimoto, T., & Inagaki, Y. (2015). Plastid genome-based phylogeny pinpointed the origin of the green-colored plastid in the dinoflagellate Lepidodinium chlorophorum. Genome Biology and Evolution, 7(4), 1133–1140. https://doi.org/10.1093/gbe/evv060

Katoh, K., & Standley, D. M. (2013). MAFFT multiple sequence alignment software version 7: improvements in performance and usability. Molecular Biology and Evolution, 30(4), 772–780. https://doi.org/10.1093/molbev/mst010

Kaundal, R., Saini, R., & Zhao, P. X. (2010). Combining Machine Learning and Homology-Based Approaches to Accurately Predict Subcellular Localization in Arabidopsis. Plant Physiology, 154(1), 36–54. https://doi.org/10.1104/pp.110.156851

Kopp, C., Pernice, M., Domart-Coulon, I., Djediat, C., Spangenberg, J. E., Alexander, D. T. L., Hignette, M., Meziane, T., Meibom, A., Orphan, V., & McFall-Ngai, M. J. (2013). Highly Dynamic Cellular-Level Response of Symbiotic Coral to a Sudden Increase in Environmental Nitrogen. mBio, 4(3), e00052–00013. https://doi.org/doi:10.1128/mBio.00052-13

Korf, I. (2004). Gene finding in novel genomes. BMC Bioinformatics, 5(1), 59. https://doi.org/10.1186/1471-2105-5-59

Krochko, J. E., Abrams, G. D., Loewen, M. K., Abrams, S. R., & Cutler, A. J. (1998). (+)-Abscisic acid 8’-hydroxylase is a cytochrome P450 monooxygenase. Plant Physiology, 118(3), 849–860. https://doi.org/10.1104/pp.118.3.849

Laetsch, D., & Blaxter, M. (2017). BlobTools: Interrogation of genome assemblies F1000Research, 6, 1287. https://doi.org/10.12688/f1000research.12232.1

LaJeunesse, T. C. (2005). “Species” radiations of symbiotic dinoflagellates in the Atlantic and Indo-Pacific since the Miocene-Pliocene transition. Molecular Biology and Evolution, 22(3), 570–581.

LaJeunesse, T. C., Parkinson, J. E., Gabrielson, P. W., Jeong, H. J., Reimer, J. D., Voolstra, C. R., & Santos, S. R. (2018). Systematic Revision of Symbiodiniaceae Highlights the Antiquity and Diversity of Coral Endosymbionts. Current Biology, 28(16), 2570-2580.e2576. https://doi.org/https://doi.org/10.1016/j.cub.2018.07.008

Langmead, B., & Salzberg, S. L. (2012). Fast gapped-read alignment with Bowtie 2. Nature Methods, 9(4), 357–359. https://doi.org/10.1038/nmeth.1923

Levin, R. A., Beltran, V. H., Hill, R., Kjelleberg, S., McDougald, D., Steinberg, P. D., & van Oppen, M. J. H. (2016). Sex, scavengers, and chaperones: transcriptome secrets of divergent Symbiodinium thermal tolerances. Molecular Biology and Evolution, 33(11), 2201–2215. https://doi.org/10.1093/molbev/msw201

Li, H. (2018). Minimap2: pairwise alignment for nucleotide sequences. Bioinformatics, 34(18), 3094–3100. https://doi.org/10.1093/bioinformatics/bty191

Li, T., Yu, L., Song, B., Song, Y., Li, L., Lin, X., & Lin, S. (2020). Genome improvement and core gene set refinement of Fugacium kawagutii. Microorganisms, 8(1), 102.

Li, W., & Godzik, A. (2006). Cd-hit: a fast program for clustering and comparing large sets of protein or nucleotide sequences. Bioinformatics, 22(13), 1658–1659. https://doi.org/10.1093/bioinformatics/btl158

Lin, S., Cheng, S., Song, B., Zhong, X., Lin, X., Li, W., Li, L., Zhang, Y., Zhang, H., & Ji, Z. (2015). The Symbiodinium kawagutii genome illuminates dinoflagellate gene expression and coral symbiosis. Science, 350(6261), 691–694.

Lin, S., Song, B., & Morse, D. (2021). Spatial organization of dinoflagellate genomes: Novel insights and remaining critical questions. Journal of Phycology, 57(6), 1674–1678.

Liu, H., Stephens, T. G., González-Pech, R. A., Beltran, V. H., Lapeyre, B., Bongaerts, P., Cooke, I., Aranda, M., Bourne, D. G., Forêt, S., Miller, D. J., van Oppen, M. J. H., Voolstra, C. R., Ragan, M. A., & Chan, C. X. (2018). Symbiodinium genomes reveal adaptive evolution of functions related to coral-dinoflagellate symbiosis. Communications Biology, 1(1), 95. https://doi.org/10.1038/s42003-018-0098-3

Lo, R., Dougan, K. E., Chen, Y., Shah, S., Bhattacharya, D., & Chan, C. X. (2022). Alignment-Free Analysis of Whole-Genome Sequences From Symbiodiniaceae Reveals Different Phylogenetic Signals in Distinct Regions [Original Research]. Frontiers in Plant Science, 13. https://doi.org/10.3389/fpls.2022.815714

Lomsadze, A., Ter-Hovhannisyan, V., Chernoff, Y. O., & Borodovsky, M. (2005). Gene identification in novel eukaryotic genomes by self-training algorithm. Nucleic Acids Research, 33(20), 6494–6506. https://doi.org/10.1093/nar/gki937

Manni, M., Berkeley, M. R., Seppey, M., Simão, F. A., & Zdobnov, E. M. (2021). BUSCO update: novel and streamlined workflows along with broader and deeper phylogenetic coverage for scoring of eukaryotic, prokaryotic, and viral genomes. Molecular Biology and Evolution, 38(10), 4647–4654.

Marçais, G., & Kingsford, C. (2011). A fast, lock-free approach for efficient parallel counting of occurrences of k-mers. Bioinformatics, 27(6), 764–770.

Marinov, G. K., Trevino, A. E., Xiang, T., Kundaje, A., Grossman, A. R., & Greenleaf, W. J. (2021). Transcription-dependent domain-scale three-dimensional genome organization in the dinoflagellate Breviolum minutum. Nature Genetics, 53(5), 613–617.

Melis, A., & Chen, H.-C. (2005). Chloroplast sulfate transport in green algae –genes, proteins and effects. Photosynthesis Research, 86(3), 299–307. https://doi.org/10.1007/s11120-005-7382-z

Minh, B. Q., Schmidt, H. A., Chernomor, O., Schrempf, D., Woodhams, M. D., von Haeseler, A., & Lanfear, R. (2020). IQ-TREE 2: New Models and Efficient Methods for Phylogenetic Inference in the Genomic Era. Molecular Biology and Evolution, 37(5), 1530–1534. https://doi.org/10.1093/molbev/msaa015

Morozov, A. A., & Galachyants, Y. P. (2019). Diatom genes originating from red and green algae: Implications for the secondary endosymbiosis models. Marine Genomics, 45, 72–78. https://doi.org/https://doi.org/10.1016/j.margen.2019.02.003

Moustafa, A., Beszteri, B., Maier, U. G., Bowler, C., Valentin, K., & Bhattacharya, D. (2009). Genomic footprints of a cryptic plastid endosymbiosis in diatoms. Science, 324(5935), 1724–1726. https://doi.org/10.1126/science.1172983

Muscatine, L., Falkowski, P. G., Porter, J. W., Dubinsky, Z., & Smith, D. C. (1984). Fate of photosynthetic fixed carbon in light-and shade-adapted colonies of the symbiotic coral Stylophora pistillata. Proceedings of the Royal Society of London. Series B. Biological Sciences, 222(1227), 181–202. https://doi.org/doi:10.1098/rspb.1984.0058

Nand, A., Zhan, Y., Salazar, O. R., Aranda, M., Voolstra, C. R., & Dekker, J. (2021). Genetic and spatial organization of the unusual chromosomes of the dinoflagellate Symbiodinium microadriaticum. Nature Genetics, 53(5), 618–629.

Nevo, Z., & Sharon, N. (1969). The cell wall of Peridinium westii, a non cellulosic glucan. Biochimica et Biophysica Acta (BBA) - Biomembranes, 173(2), 161–175. https://doi.org/https://doi.org/10.1016/0005-2736(69)90099-6

Price, D. C., & Bhattacharya, D. (2017). Robust Dinoflagellata phylogeny inferred from public transcriptome databases. Journal of Phycology, 53(3), 725–729.

Ranallo-Benavidez, T. R., Jaron, K. S., & Schatz, M. C. (2020). GenomeScope 2.0 and Smudgeplot for reference-free profiling of polyploid genomes. Nature Communications, 11(1), 1–10.

Remmert, M., Biegert, A., Hauser, A., & Söding, J. (2012). HHblits: lightning-fast iterative protein sequence searching by HMM-HMM alignment. Nature Methods, 9(2), 173–175. https://doi.org/10.1038/nmeth.1818

Rosic, N. N., & Hoegh-Guldberg, O. (2010). A method for extracting a high-quality RNA from Symbiodinium sp. Journal of Applied Phycology, 22(2), 139–146. https://doi.org/10.1007/s10811-009-9433-x

Rosic, N. N., Pernice, M., Dunn, S., Dove, S., & Hoegh-Guldberg, O. (2010). Differential regulation by heat stress of novel cytochrome P450 genes from the dinoflagellate symbionts of reef-building corals. Applied and Environmental Microbiology, 76(9), 2823–2829. https://doi.org/10.1128/AEM.02984-09

Salmeán, A. A., Duffieux, D., Harholt, J., Qin, F., Michel, G., Czjzek, M., Willats, W. G. T., & Hervé, C. (2017). Insoluble (1 → 3), (1 → 4)-β-D-glucan is a component of cell walls in brown algae (Phaeophyceae) and is masked by alginates in tissues. Scientific Reports, 7(1), 2880. https://doi.org/10.1038/s41598-017-03081-5

Sarai, C., Tanifuji, G., Nakayama, T., Kamikawa, R., Takahashi, K., Yazaki, E., Matsuo, E., Miyashita, H., Ishida, K.-i., Iwataki, M., & Inagaki, Y. (2020). Dinoflagellates with relic endosymbiont nuclei as models for elucidating organellogenesis. Proceedings of the National Academy of Sciences, 117(10), 5364–5375. https://doi.org/doi:10.1073/pnas.1911884117

Shoguchi, E., Beedessee, G., Tada, I., Hisata, K., Kawashima, T., Takeuchi, T., Arakaki, N., Fujie, M., Koyanagi, R., & Roy, M. C. (2018). Two divergent Symbiodinium genomes reveal conservation of a gene cluster for sunscreen biosynthesis and recently lost genes. BMC Genomics, 19(1), 1–11.

Stanke, M., Keller, O., Gunduz, I., Hayes, A., Waack, S., & Morgenstern, B. (2006). AUGUSTUS: ab initio prediction of alternative transcripts. Nucleic Acids Research, 34(suppl_2), W435–W439. https://doi.org/10.1093/nar/gkl200

Stephens, T. G., Bhattacharya, D., Ragan, M. A., & Chan, C. X. (2016). PhySortR: a fast, flexible tool for sorting phylogenetic trees in R. PeerJ, 4, e2038.

Stephens, T. G., González-Pech, R. A., Cheng, Y., Mohamed, A. R., Burt, D. W., Bhattacharya, D., Ragan, M. A., & Chan, C. X. (2020). Genomes of the dinoflagellate Polarella glacialis encode tandemly repeated single-exon genes with adaptive functions. BMC Biology, 18(1), 56. https://doi.org/10.1186/s12915-020-00782-8

Stephens, T. G., Ragan, M. A., Bhattacharya, D., & Chan, C. X. (2018). Core genes in diverse dinoflagellate lineages include a wealth of conserved dark genes with unknown functions. Scientific Reports, 8(1), 1–11.

Wang, Y., Li, J., & Paterson, A. H. (2013). MCScanX-transposed: detecting transposed gene duplications based on multiple colinearity scans. Bioinformatics, 29(11), 1458–1460.

Wilson, K., Li, Y., Whan, V., Lehnert, S., Byrne, K., Moore, S., Pongsomboon, S., Tassanakajon, A., Rosenberg, G., Ballment, E., Fayazi, Z., Swan, J., Kenway, M., & Benzie, J. (2002). Genetic mapping of the black tiger shrimp Penaeus monodon with amplified fragment length polymorphism. Aquaculture, 204(3), 297–309. https://doi.org/https://doi.org/10.1016/S0044-8486(01)00842-0

Yoon, H. S., Hackett, J. D., & Bhattacharya, D. (2002). A single origin of the peridinin-and fucoxanthin-containing plastids in dinoflagellates through tertiary endosymbiosis. Proceedings of the National Academy of Sciences, 99(18), 11724–11729. https://doi.org/doi:10.1073/pnas.172234799

Yoon, H. S., Hackett, J. D., Van Dolah, F. M., Nosenko, T., Lidie, K. L., & Bhattacharya, D. (2005). Tertiary endosymbiosis driven genome evolution in dinoflagellate algae. Molecular Biology and Evolution, 22(5), 1299–1308.

Zimin, A. V., Puiu, D., Luo, M.-C., Zhu, T., Koren, S., Marçais, G., Yorke, J. A., Dvořák, J., & Salzberg, S. L. (2017). Hybrid assembly of the large and highly repetitive genome of Aegilops tauschii, a progenitor of bread wheat, with the MaSuRCA mega-reads algorithm. Genome Research, 27(5), 787–792. https://doi.org/10.1101/gr.213405.116

