## Supplementary Figures for "Improved *Cladocopium goreaui* genome assembly reveals features of a facultative coral symbiont and the complex evolutionary history of dinoflagellate genes"

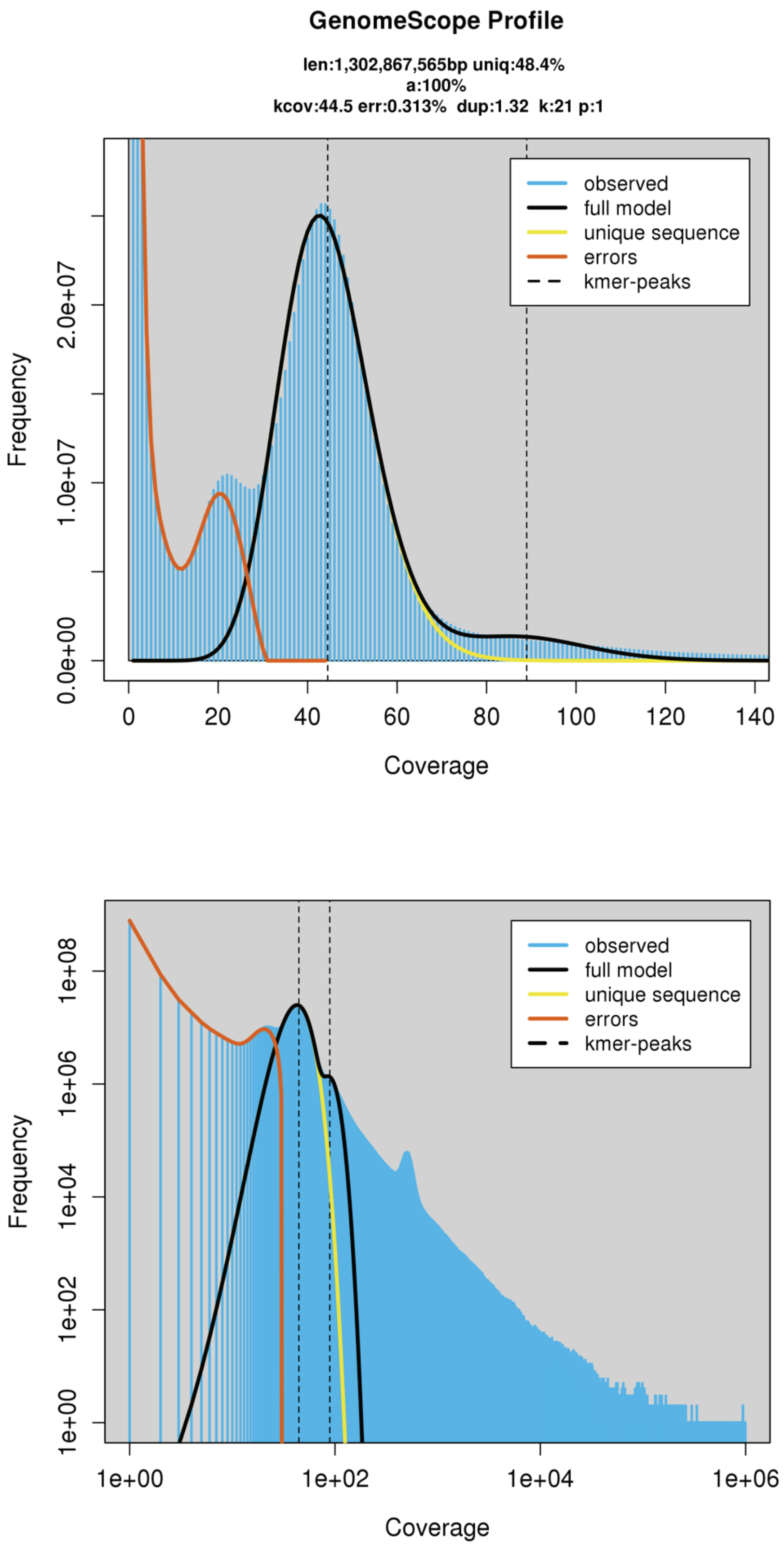


**Figure S1.** Genome size estimation for *Cladocopium goreaui* using GenomeScope v2.0.

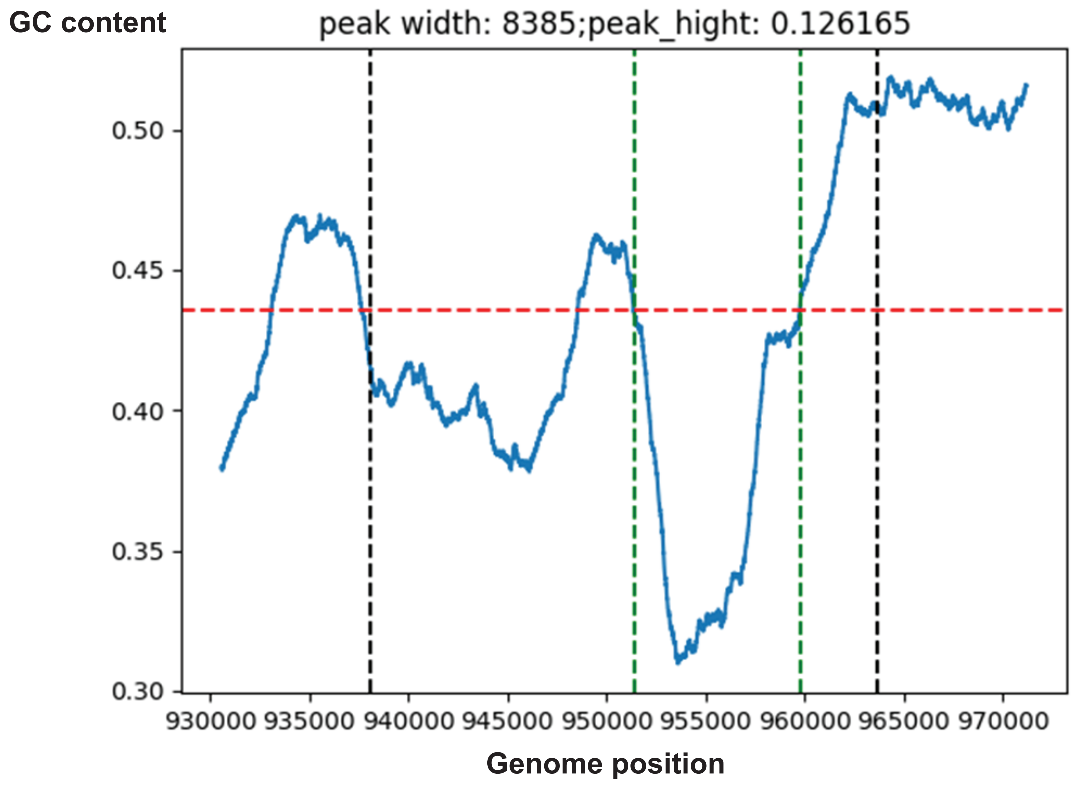


**Figure S2.** An example of G+C dip observed in the *C. goreaui* genome of a putative boundary of topologically associated domain (TAD).

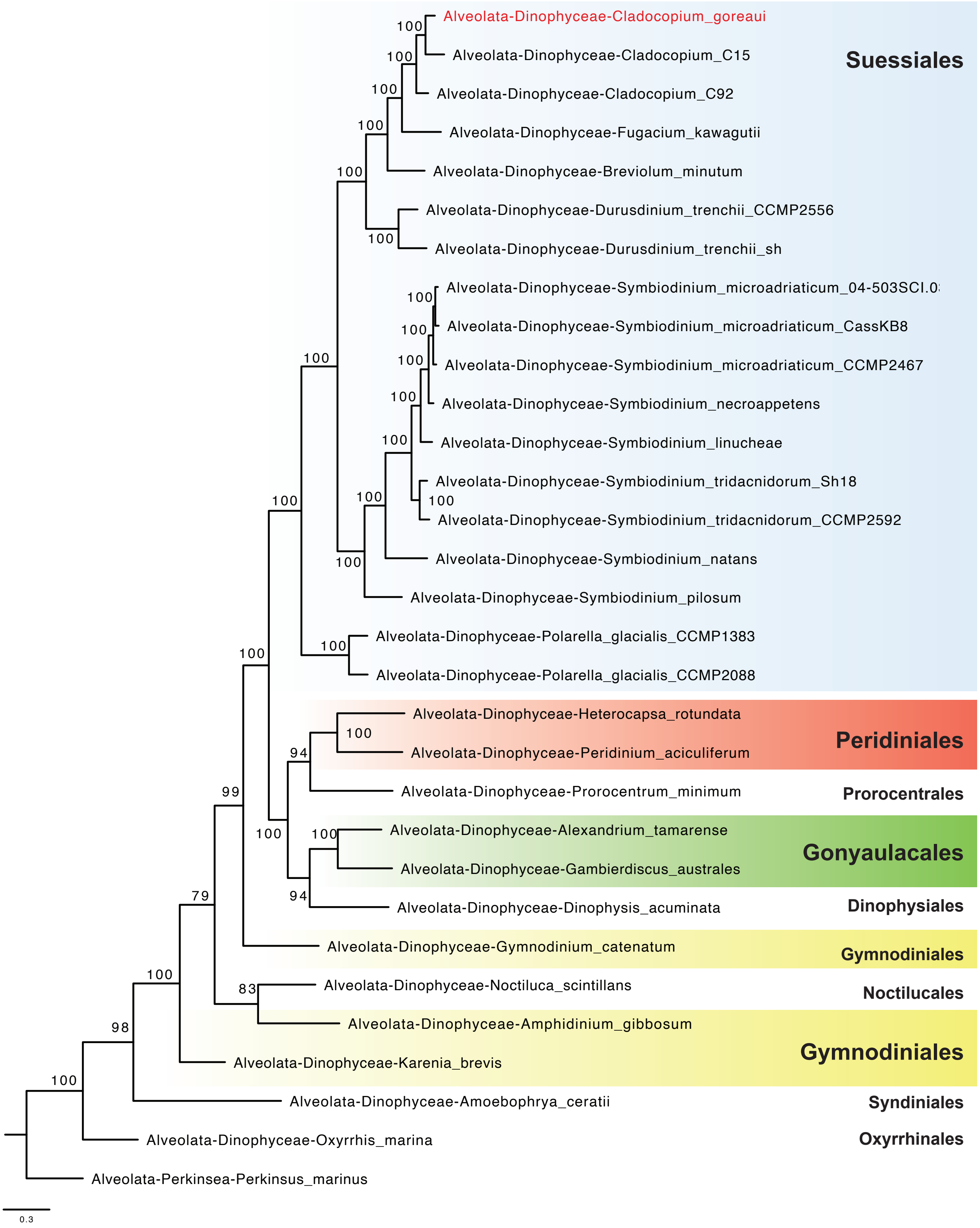


**Figure S3.** Species tree of 30 dinoflagellate taxa and *Perkinsus* as outgroup, inferred from 3,411 strictly orthologous (single-copy) protein sets.

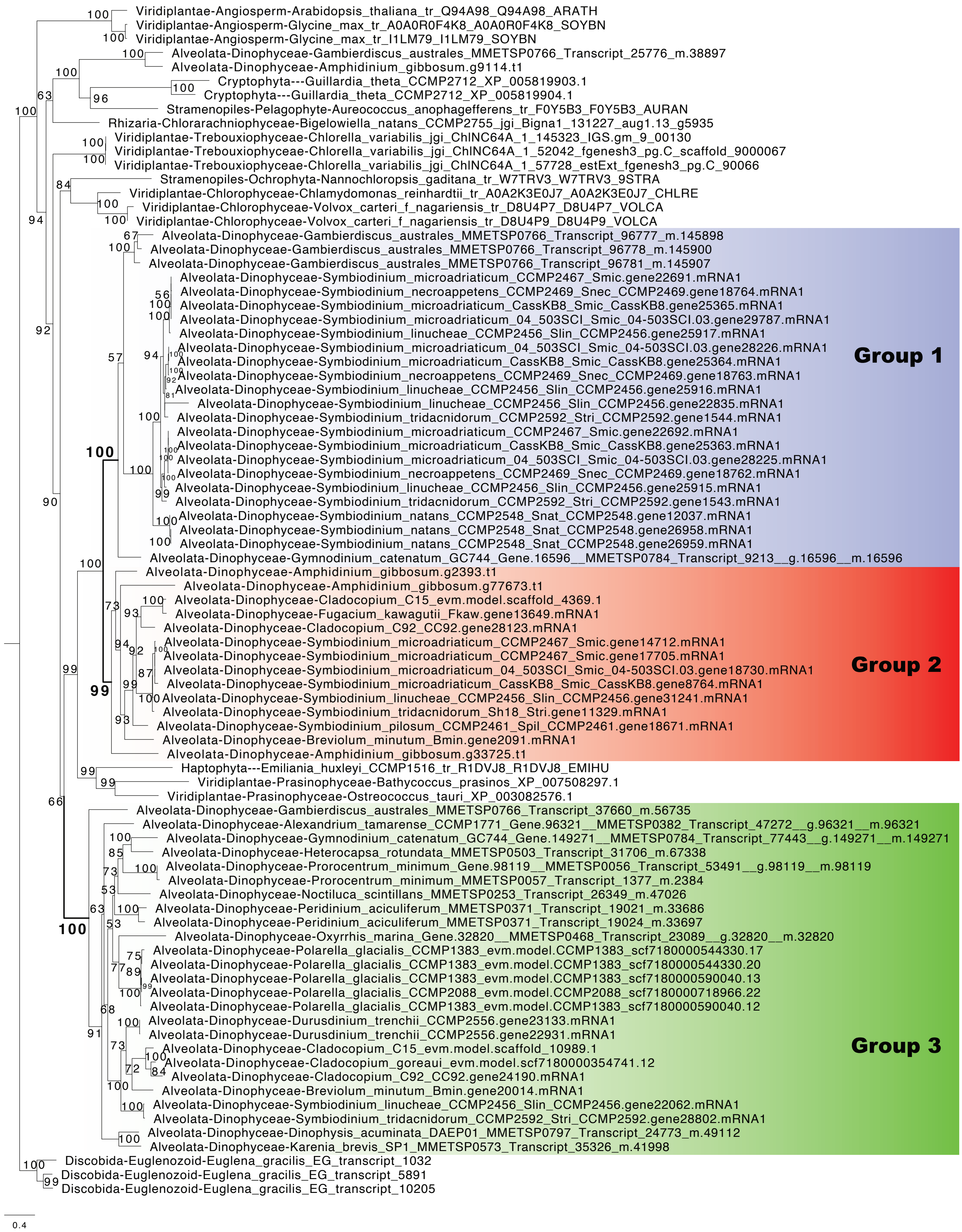


**Figure S4.** Maximum likelihood tree showing gene expansion of a green algal derived gene family.

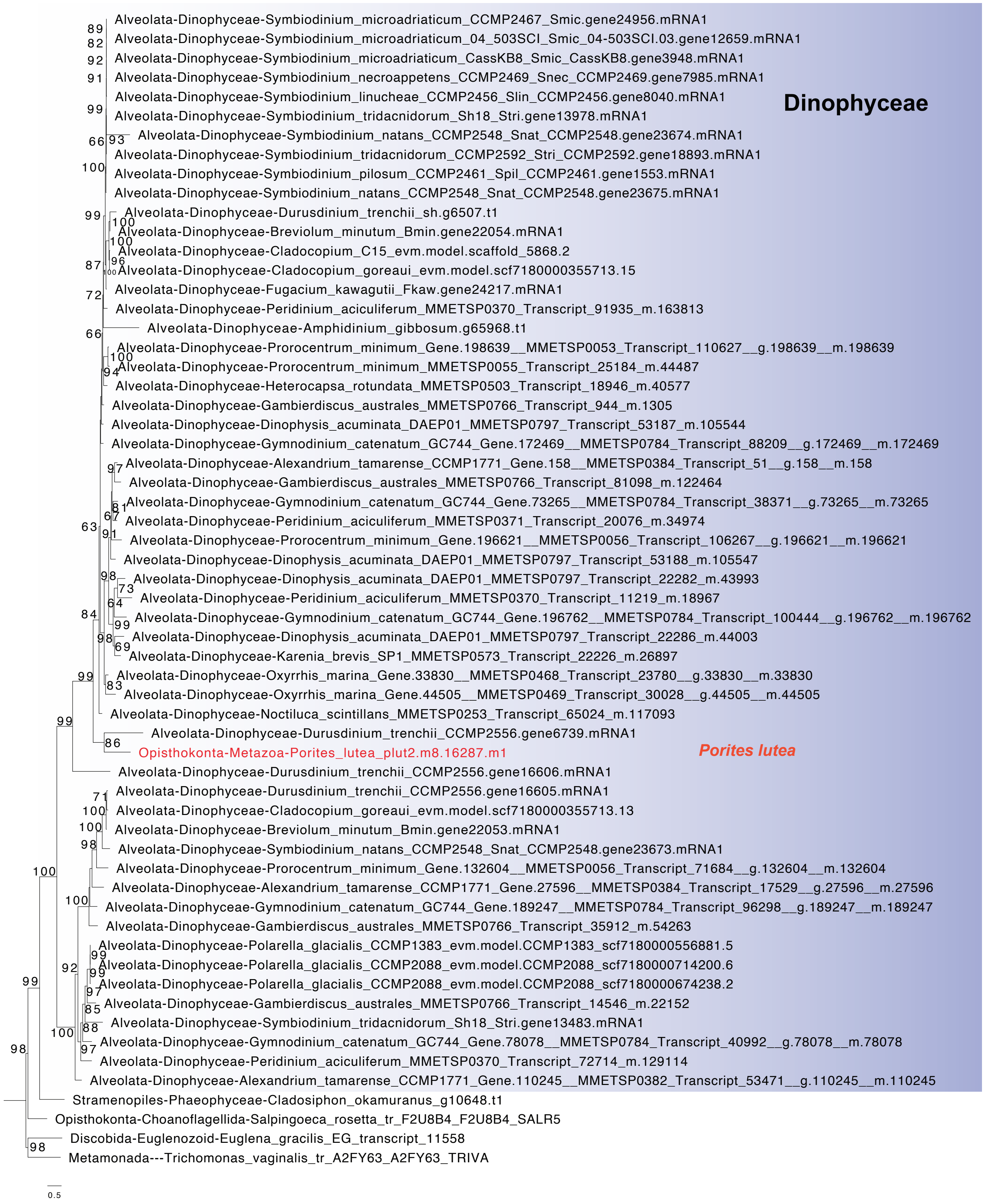


**Figure S5.** Maximum likelihood tree of phosphatidylinositol 4-phosphate 5-kinase showing possible misidentification of the sequence from the dinoflagellate symbiont associated with the coral.

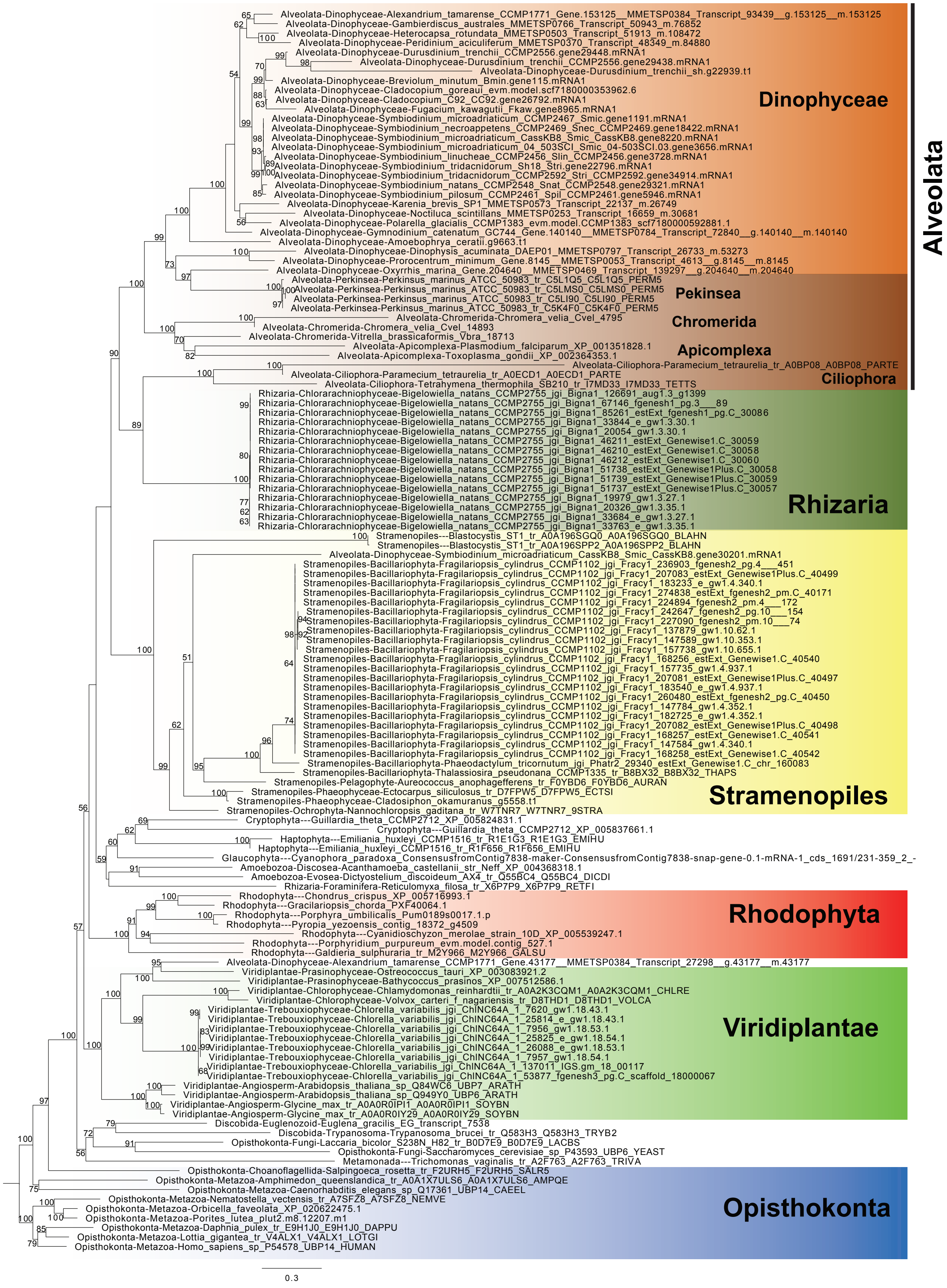


**Figure S6.** Maximum likelihood tree of ubiquitin carboxyl-terminal hydrolase showing strong evidence of vertical inheritance

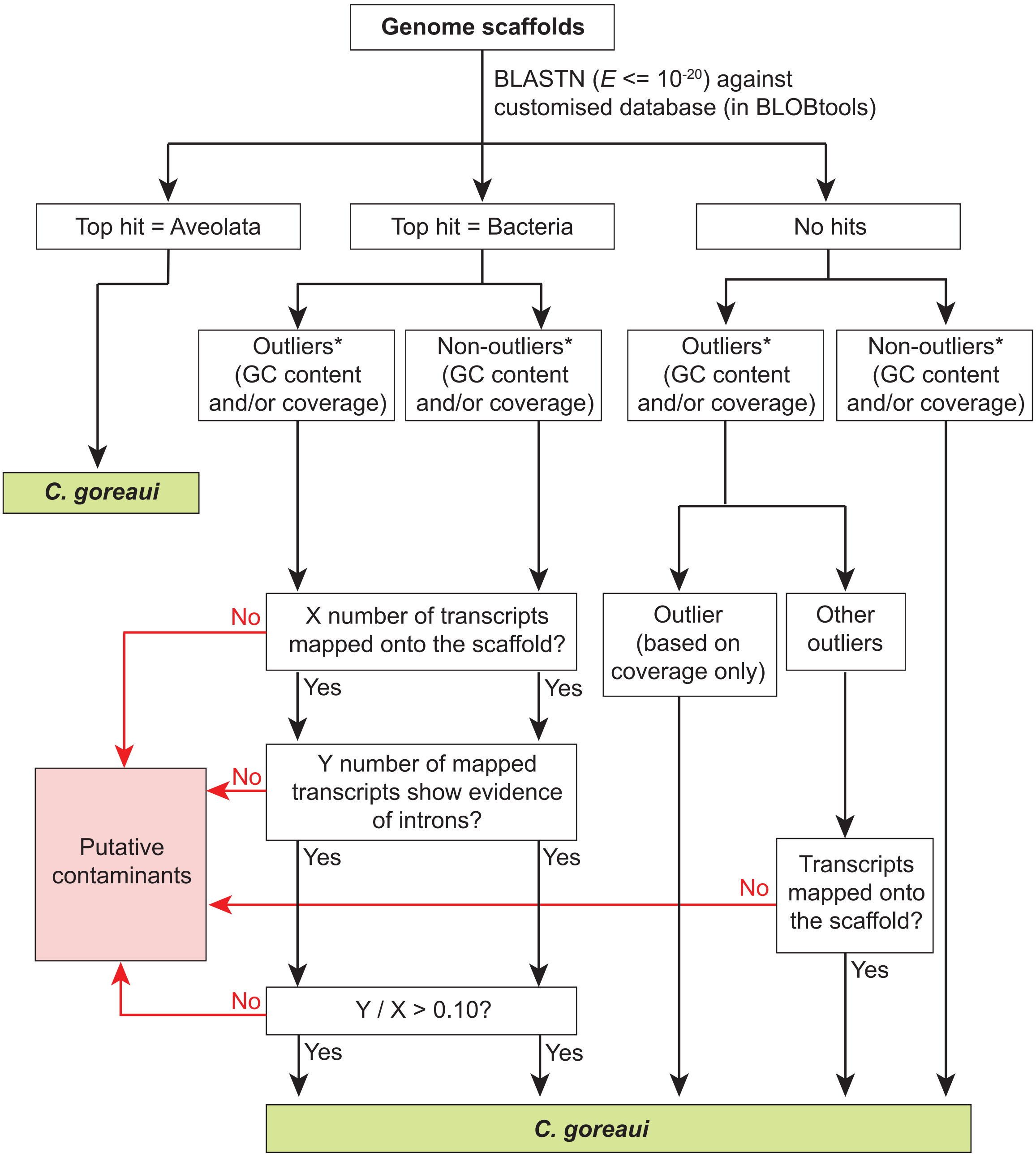


*: outliers are determined using BLOBtools. Scaffolds for which G+C content and/or read coverage is external to the range of median ± 1.5 × interquartile range (IQR) are considered as outliers.

**Figure S7.** Decision tree for identification and removal of putative contaminant sequences.
